# Mechanism for differential recruitment of orbitostriatal transmission during outcomes and actions in alcohol dependence

**DOI:** 10.1101/2020.12.04.411819

**Authors:** Rafael Renteria, Christian Cazares, Emily T. Baltz, Drew C. Schreiner, Ege A. Yalcinbas, Thomas Steinkellner, Thomas S. Hnasko, Christina M. Gremel

## Abstract

Psychiatric disease often produces symptoms that have divergent effects on neural activity. For example, in drug dependence, dysfunctional value-based decision-making and compulsive-like actions have been linked to hypo- and hyper-activity of orbital frontal cortex (OFC)-basal ganglia circuits, respectively, however, the underlying mechanisms are unknown. Here we show that alcohol dependence enhanced activity in OFC terminals in dorsal striatum (OFC-DS) associated with actions, but reduced activity of the same terminals during periods of outcome retrieval, corresponding with a loss of outcome control over decision-making. Disrupted OFC-DS terminal activity was due to a dysfunction of dopamine-type 1 receptors on spiny projection neurons (D1R SPNs) that resulted in increased retrograde endocannabinoid (eCB) signaling at OFC-D1R SPN synapses reducing OFC-DS transmission. Blocking CB1 receptors restored OFC-DS activity *in vivo* and rescued outcome-based control over decision-making. These findings demonstrate a circuit-, synapse-, and computation specific mechanism gating OFC activity following the induction of alcohol dependence.

## Introduction

The DSM-5 categorizes alcohol as one of the few drugs whose dependence can produce neurocognitive disorders (American Psychiatric Association, 2013). Thus, treatment of alcohol dependence would benefit from understanding mechanisms producing impaired cognition. Value-based decision-making is one such cognitive process, with its disruption contributing to poor daily function, that when combined with the emergence of compulsive control contributes to excessive drug-seeking, and relapse (Belin et al., 2013; Everitt and Robbins, 2016; Gremel and Lovinger, 2017; Hogarth et al., 2013; Lüscher et al., 2020). Recently, there has been renewed focus on orbital frontal cortex (OFC) circuits in addiction and addiction-like behaviors (Everitt and Robbins, 2016; Lüscher et al., 2020; Moorman, 2018; Schoenbaum et al., 2016, 2006), as they play a key role in computations related to decision-making and the evaluation of outcomes (Fellows, 2007; Stalnaker et al., 2015; Wallis, 2007). However, a mechanistic understanding of how drug-dependence disrupts the ability of OFC and its downstream circuits to represent information necessary for appropriate control over decision-making is lacking.

Indeed, drug dependence in general has been widely associated with reduced OFC activity and output (Catafau et al., 1999; Lüscher et al., 2020; Schoenbaum et al., 2016, 2006; Volkow et al., 1997; Volkow and Fowler, 1994). In alcohol dependent individuals, reduced OFC activity correlates with aberrant value-based decision-making (Boettiger et al., 2007; Duka et al., 2011; Forbes et al., 2014). However, OFC hyperactivity has been observed in alcohol dependence in response to predictive information (Hermann et al., 2006; Reinhard et al., 2015; Tapert et al., 2003) and during approach behaviors (Ernst et al., 2014). In concert, both rodent and non-human primate models of alcohol dependence have identified long-lasting changes to OFC neuron excitability, structure, and transmission (Nimitvilai et al., 2017, 2016; Renteria et al., 2018). Thus, it seems likely that alcohol dependence differentially alters the contribution of OFC circuits and their downstream controllers to varied decision-making processes, thereby enhancing some computations while reducing others, leading to alterations in behavioral control.

One such circuit is OFC excitatory projections into basal ganglia, with synapses onto direct and indirect pathways in the dorsal striatum (DS) (Renteria et al., 2018; Wall et al., 2013). Excitatory transmission from OFC-DS projections carries information that may support value-based decision-making, as inhibition of transmission results in habitual control (Gremel et al., 2016). We recently found that alcohol dependence selectively attenuates output of this circuit, with a long-lasting reduction in OFC-DS transmission onto dopamine-type 1 (D1) spiny projection neurons (SPNs) of the direct pathway (Renteria et al., 2018). In addition, alcohol dependence produces a loss of value-based decision-making (Corbit et al., 2012; Dickinson et al., 2002; Renteria et al., 2018) and reductions in associated OFC activity (Catafau et al., 1999; Lüscher et al., 2020; Schoenbaum et al., 2016, 2006; Volkow et al., 1997; Volkow and Fowler, 1994). Indeed, this behavioral phenotype was rescued by increasing OFC activity (Renteria et al., 2018). Together, these findings suggest that dependence may result in a reduced communication of value-related information from OFC into basal-ganglia circuits to support decision-making.

However, dependence has also been associated with compulsive phenotypes (Everitt and Robbins, 2016, 2005; Lüscher et al., 2020) and hyperactivity of OFC-DS circuits has been implicated in obsessive compulsive disorder (OCD) (Lüscher et al., 2020; Milad and Rauch, 2012; Pauls et al., 2014; Robbins et al., 2019), as well as animal models of compulsive (Ahmari et al., 2013) and compulsive addiction-like behaviors (Pascoli et al., 2018, 2015). Indeed, increased OFC terminal activity was observed in mice when they self-stimulated ventral tegmental area (VTA) dopamine neurons, despite the presence of a punishing foot-shock (Pascoli et al., 2018). This corresponded to post-synaptic potentiation of OFC-DS transmission (Pascoli et al., 2018), suggesting that increased transmission from OFC into basal ganglia circuits contributes to the emergence of compulsive actions. Thus, OFC-DS circuits are poised to regulate both compulsive action and value information, computations that are both implicated in addiction and may be differentially altered in psychiatric disease.

Here we examine mechanisms underlying how drug dependence affects OFC-DS circuit information representation and its control over decision-making. Using a well-established model of alcohol dependence, we find that dependence selectively enhances OFC-DS activity associated with action while reducing OFC-DS activity during periods associated with outcome retrieval through down-regulation of D1 receptor function in SPNs and enhanced endocannabinoid (eCB) signaling at OFC-D1 SPN synapses in the DS. Restoring OFC-DS activity rescues value-based decision-making. Our data has important implications for hypotheses regarding compulsive and habitual phenotypes observed in addiction.

## Results

### Altered encoding of OFC-DS terminal activity following alcohol dependence

To examine whether alcohol dependence alters information sent to dorsal striatum, we used fiber photometry to measure calcium activity of OFC terminals in DS during self-initiated value-based decision-making. We utilized a well-validated and commonly used model of alcohol dependence, chronic intermittent ethanol (CIE) vapor exposure (Becker, 1994; Becker and Lopez, 2004; Griffin 3rd et al., 2009; Lopez and Becker, 2005) (Figure 1A). Performed in 3 cohort replications, mice (Air *n* = 10, CIE *n* = 12) were exposed to ethanol vapor for 16 hours each day for 4 consecutive days followed by a 3-day withdrawal. This cycle of ethanol vapor and withdrawal was repeated for 4 consecutive weeks and resulted in blood ethanol concentrations of 33.70 ± 2.24 mM (mean ± SEM) (See Methods). Prior to CIE procedures, mice were injected with AAV-hSynapsin1-axon-GCaMP6s-P2A-mRuby3 in the OFC and implanted with a fiber targeted to the medial DS to record OFC terminal activity (See Methods) (Figure 1B-D). Alcohol withdrawal has been delineated into two differing stages, an acute withdrawal period lasting 2-3 days, and longer protracted withdrawal period that can last 3 or more months (American Psychiatric Association, 2013; Heilig et al., 2010). To avoid effects of acute withdrawal and examine potential alterations in the protracted withdrawal period, mice were food restricted and five days after the last vapor exposure were trained to lever press for a food reward under a random ratio schedule of reinforcement to bias value-based decision-making, measured as a sensitivity to outcome devaluation (Dickinson et al., 1983; Gremel and Costa, 2013) (Figure 1E).

**Figure 1.**
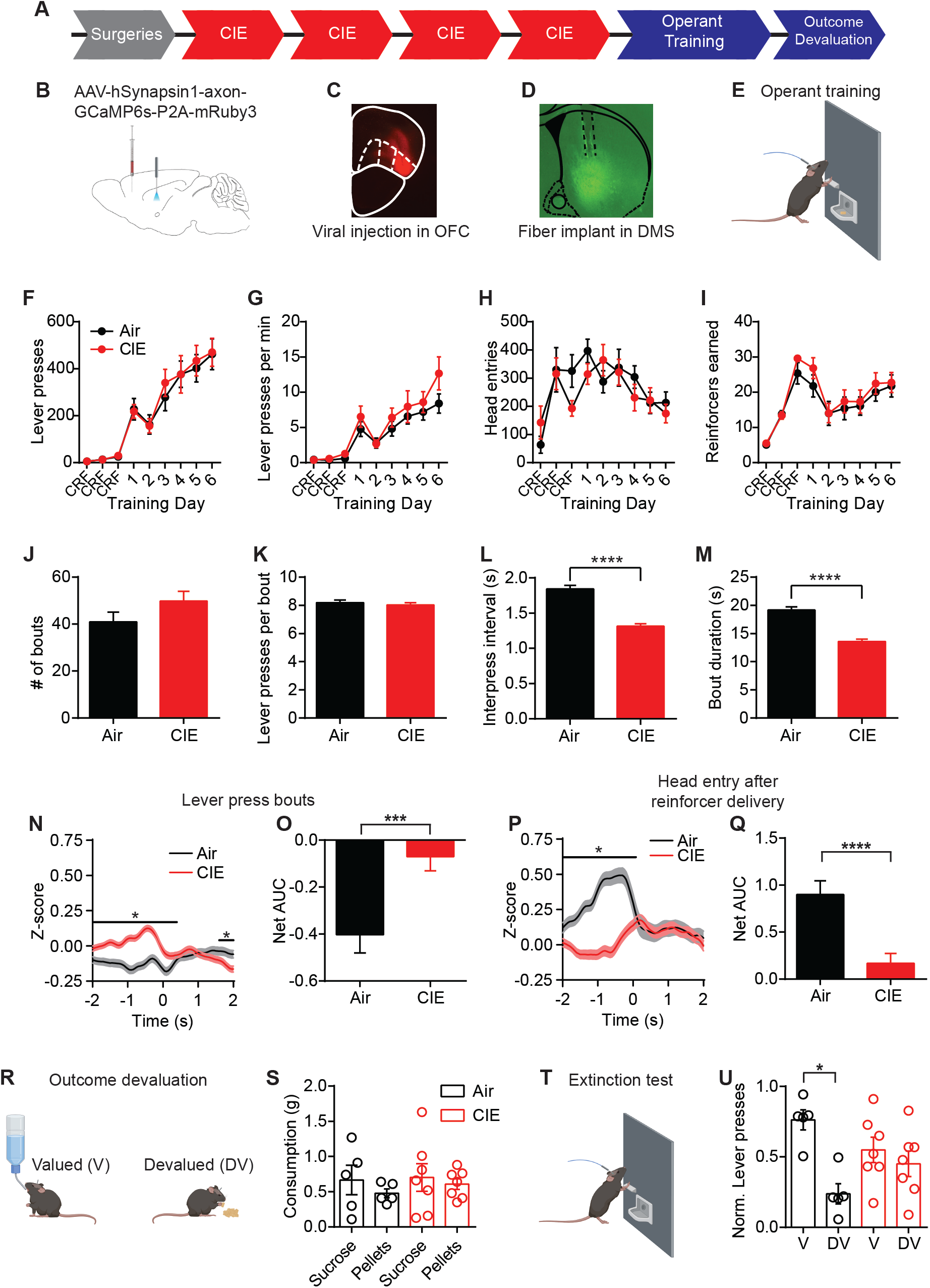
CIE induced alterations in OFC terminal activity in the DS during instrumental responding. **(A)** Experimental timeline that includes surgeries followed by 4 cycles of CIE exposure, operant training and outcome devaluation. **(B)** Schematic of viral injections in the OFC and fiber implant in the DS. **(C)** Example viral expression in OFC. **(D)** Example OFC terminal expression and fiber placement in the DS. **(E)** Lever press training for a food pellet under random ratio schedule of reinforcement. **(F)** Lever presses, **(G)** response rate, **(H)** head entries and **(I)** reinforcers earned during training in Air (*n* = 10 mice, 3 vapor cohorts) and CIE (*n* = 12 mice, 3 vapor cohorts) mice. **(J)** Average number of lever press bouts in Air and CIE mice. **(K)** Average number of lever presses per bout. **(L)** Average inter-press interval within lever press bouts. **(M)** Average bout duration. **(N)** Z-score of ΔF/F GCaMP6s traces recorded in OFC terminals during lever press bouts. **(O)** Net area under the curve (AUC) of Z-score GCaMP6s signal aligned to the start of a lever press bout. **(P)** Z-score of ΔF/F GCaMP6s trace during the first head entry after reinforcer delivery. **(Q)** Net AUC of Z-score GCaMP6s signal during first head entry after reinforcer delivery. **(R)** Schematic of outcome devaluation procedure. **(S)** Consumption of sucrose and pellets in Air (*n* = 5 mice) and CIE (*n* = 7 mice) mice during one-hour free feed period. **(T)** Schematic of extinction test following free feed period. **(U)** Distribution of lever presses across valued (V) and devalued (DV) days. Data points and bar graphs represent mean ± SEM. Bonferroni-corrected post-hoc test of repeated-measures ANOVA, two-sided FDR-corrrected permutation test, or KS-test, *p < 0.05, ****p < 0.0001.

Alcohol dependence did not prevent mice from learning to lever press for food. Air control mice and CIE exposed mice made the same number of lever presses (repeated-measures ANOVA: main effect of Training day: F_(8, 160)_ = 43.10, p < 0.0001; no main effect of CIE exposure or interaction: Fs<0.16) (Figure 1F), and pressed the lever at roughly equal rates (repeated-measures ANOVA: main effect of Training day: F_(8, 160)_ = 27.42, p < 0.0001; no main effect of treatment or interaction: Fs < 1.22) (Figure 1G). The two groups made similar number of head entries across training (repeated-measures ANOVA: main effect of Training day: F_(8, 160)_ = 5.92, p<0.0001; no main effect of CIE exposure or interaction: Fs < 1.29) (Figure 1H), and earned similar numbers of rewards (repeated-measures ANOVA: main effect of Training day: F_(8, 160)_ = 19.42, p<0.0001; no main effect of treatment or interaction: Fs < 0.60) (Figure 1I). As mice self-initiate, self-terminate their own behavior, it appeared they often made multiple lever presses in close succession. We used bout analyses to examine whether mice organized their lever presses into distinguishable patterns (See Methods). Mice grouped their lever presses into distinct bouts, and Air and CIE mice had similar numbers of bouts (Air: 40.92 ± 4.15, *n* = 37; CIE: 49.78 ± 4.17, *n* = 37; non-parametric two-sample Kolmogorov-Smirnov (KS) test: D = 0.30, p = 0.07) (Figure 1J) with similar numbers of lever presses per bout (Air: 8.19 ± 0.18, *n* = 1514; CIE: 8.032 ± 0.16, *n* = 1842; KS-test: D = 0.04, p = 0.14) (Figure 1K). However, Air and CIE mice differentially executed these bouts, with CIE mice having shorter intervals between lever presses within a bout (Inter-press Interval, Air: 1.84 s ± 0.05, *n* = 1514; CIE: 1.31 s ± 0.04, *n* = 1842; KS-test: D = 0.17, p < 0.0001) (Figure 1L) resulting in shorter bout durations (Air: 19.15 s ± 0.59, *n* = 1514; CIE: 13.59 s ± 0.41, *n* = 1842; KS-test: D = 0.14, p < 0.0001) (Figure 1M). Thus, the prior induction of alcohol dependence led to a change in action execution, manifested as faster completion of lever press bouts.

When we examined OFC-DS Ca^2+^ activity during decision-making, we observed modulation of calcium transients during action epochs (−2s to 2s after the onset of a lever press bout) and outcome retrieval epochs (−2s to 2s around head-entry into food receptacle following reward delivery) during decision-making. As mice can readily see the food reward prior to making a head entry, this outcome retrieval period encompasses reward perception, retrieval and potentially consumption. We examined whether modulations of calcium transients were differentially affected by alcohol dependence. First, we saw CIE led to increased OFC-DS Ca^2+^ activity prior to the onset of a lever press bout (False Discovery Rate (FDR)-corrected two-sided permutation test, Air: *n* = 1363, CIE *n* = 1650, See Methods) (Figure 1N). In addition, there was a significant difference between groups in the net area under the curve (AUC) (Air: −0.40 ± 0.08, *n* = 1363; CIE: −0.05 ± 0.06, *n* = 1650; unpaired t test: t_3011_ = 3.70, p < 0.001) (Figure 1O). When we examined lever presses independent of bout grouping, we observed a similar pattern of Ca^2+^ activity in CIE mice (Figure 1-figure supplement 1). In contrast, when we examined OFC-DS activity during the period associated with outcome retrieval, following reward delivery (first head entry after reinforcer delivery), we found the opposite pattern. There, Air controls showed a large increase in OFC terminal activity, whereas in CIE mice this was significantly blunted (FDR-corrected two-sided permutation test, Air: *n* = 521, CIE: *n* = 682) (Figure 1P) and an overall decrease in the net AUC (Air: 1.00 ± 0.16, *n* = 521; CIE: 0.24 ± 0.11, *n* = 682; unpaired t test: t_1201_ = 4.12, p < 0.0001) (Figure 1Q). The above analyses were performed on lever presses collapsed across animals to maintain the variability observed in individual mouse data. We also found the same results when we compared animal session average data between groups (Figure 1-figure supplement 1), suggesting the observed differences across decision-making epochs are not driven by data from a single animal, but indeed are prevalent across animals. Together, these findings suggest that alcohol dependence leads to slightly enhanced OFC-DS terminal activity associated with actions, but substantially less OFC-DS terminal activity during the period associated with outcome retrieval.

Given the necessity of OFC-DS pathway function for successful outcome devaluation (Gremel et al., 2016), the substantial reduction in OFC-DS activity observed in CIE during periods associated with outcome retrieval suggests value-based decision-making may be impaired in these CIE mice. Thus, we used sensory-specific satiety to induce outcome devaluation (Figure 1R-U, see Methods), where reductions in lever pressing on the devalued compared to valued day are indicative of outcome devaluation. Pre-planned comparisons on normalized data to account for differences in response rates between individual mice, showed that Air reduced responding following outcome devaluation, while CIE mice did less so (repeated-measures ANOVA: main effect of valuation state: F_(1, 10)_ = 6.36, p < 0.05; no main effect of CIE exposure or interaction: Fs < 2.98; pre-planned corrected comparisons between Valued and Devalued days: Air p < 0.02; CIE p > 0.6) (Figure 1U, Figure 1-figure supplement 1). Indeed, Air mice differentially distributed lever pressing between devalued and valued days, and CIE mice did not (one-sample paired t-test vs. 0.5; Air-t_4_ = 3.68, p < 0.05; CIE t_6_ = 0.54, p = 0.30) As previous data found enhancing OFC activity was sufficient to rescue the deficits in outcome devaluation in CIE mice (Renteria et al., 2018), the current data suggest that their inability to update outcome value following devaluation may be a result of the dampened OFC-DS activity during outcome retrieval epochs (Figure 1P, Q).

### CIE does not alter CB1 receptor function or eCB long-term plasticity

The reduced calcium activity in OFC-DS terminals is suggestive of a decrease in presynaptic transmission. In dorsal striatum, OFC terminals synapse onto both direct and indirect output pathways of the basal ganglia (Wall et al., 2013). However, we previously found CIE results in a selective and long-lasting decrease in presynaptic OFC transmission onto D1 SPNs of the direct pathway, but not dopamine type-2 (D2) SPNs of the indirect pathway (Renteria et al., 2018). This reduction in OFC-D1 SPN transmission was stable and persistent across 3 weeks of chronic withdrawal (Renteria et al., 2018). As an OFC-DS projection neuron most likely sends collaterals to both D1 SPNs and D2 SPNs (Wall et al., 2013), the cell-type specific decrease in presynaptic transmission may be mediated by a CIE-induced alteration in eCB signaling (Henderson-Redmond et al., 2016; Parsons and Hurd, 2015; Pava and Woodward, 2012). eCBs are produced and released from the postsynaptic cell and act as a retrograde signal to activate presynaptic cannabinoid type 1 (CB1) receptors, resulting in an inhibition of presynaptic calcium and a subsequent decrease in neurotransmitter release (Castillo et al., 2012; Gerdeman et al., 2002; Heifets and Castillo, 2009; Kano et al., 2009; Kreitzer and Regehr, 2001; Lovinger, 2008). In the dorsal striatum, CB1 receptors have been implicated in habitual action control (Hilário et al., 2007; Nazzaro et al., 2012) and activation of CB1 receptors on OFC terminals in DS has been shown to disrupt sensitivity to changes in expected outcome value (Gremel et al., 2016).

To identify CIE-induced mechanisms by which OFC transmission is selectively decreased to D1 SPNs, nine cohort replicates of mice were exposed to 4 rounds of CIE as previously described (Figure 1A) and electrophysiological recordings were performed within 3-21 days of withdrawal (Figure 2A). Prior to CIE, D1-tdTomato or D1DR-Cre mice were injected with AAV5-CamKIIa-GFP-Cre and a Cre-dependent channelrhodopsin (ChR2) (AAV5-Ef1a-DIO-ChR2-eYFP; UNC viral vector core) in OFC to limit ChR2 expression to CamKIIa expressing excitatory projection neurons (Gremel et al., 2016; Gremel and Costa, 2013; Renteria et al., 2018; Tye et al., 2011) (Figure 2B, C). D1DR-Cre mice received an additional injection of AAV5-Flex-tdTomato in the DS to identify D1-expressing SPNs (Figure 2D). Whole-cell patch clamp recordings were performed on tdTomato expressing D1 SPNs in the DS and optically evoked EPSCs were recorded by activating ChR2 expressing OFC terminals with 470 nm light in the DS (Figure 2B).

**Figure 2.**
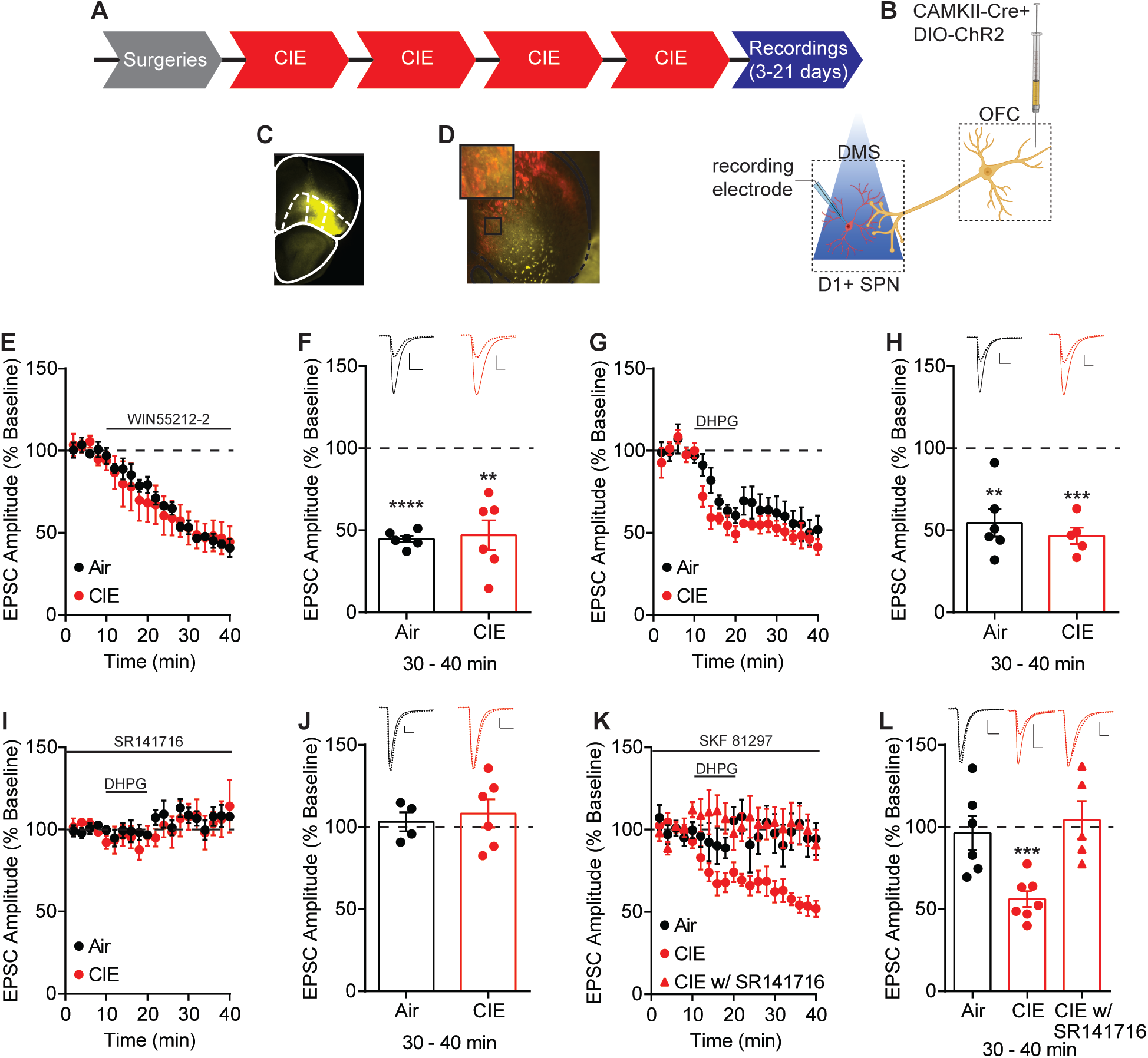
Endocannabinoid mediated plasticity of OFC transmission to D1 SPNs in Air and CIE mice. **(A)** Experimental timeline that includes surgeries followed by 4 cycles of CIE exposure and whole-cell recordings 3-21 days in withdrawal. **(B)** Schematic of viral injections in OFC and optical stimulation of OFC terminals during whole-cell recordings of D1 SPNs in the DS. **(C)** Representative viral expression of ChR2 in OFC **(D)** Example ChR2 expression of OFC terminals and expression of tdTomato in D1 SPNs in the DS. **(E)** Bath application of a CB1 receptor agonist, WIN55212-2 (1μM), during optical stimulation of OFC terminals to D1 SPNs in Air (*n* = 6 cells, 4 mice) and CIE (*n* = 6 cells, 3 mice). **(F)** Bar graphs representing the percentage change from baseline (min 0–10) after bath application of WIN55212-2 (min 30–40). **(G)** Endocannabinoid mediated long-term depression induced by bath application of a group 1 mGluR agonist, DHPG (50 μM) paired with depolarization to 0 mV in Air (*n* = 6 cells, 3 mice) and CIE (*n* = 5 cells, 3 mice).**(H)** Bar graphs representing the percentage change from baseline after bath application of DHPG. **(I)** DHPG-LTD of OFC transmission is blocked by a CB1 receptor antagonist, SR141716 (1 μM) in both Air (*n* = 4 cells, 3 mice) and CIE (*n* = 6 cells, 3 mice). **(J)** Bar graphs representing the percentage change from baseline after bath application of DHPG in the presence of SR141716. **(K)** D1 agonist (3 μM SKF 81297) blocks the expression of DHPG-LTD in Air mice (*n* = 6 cells, 4 mice) but has no effect in mice exposed to CIE (*n* = 7 cells, 5 mice; w/ SR141716 *n* = 5 cells, 2 mice). **(L)** Bar graphs representing the percentage change from baseline after bath application of DHPG in the presence of SKF 81297 or SKF 81297 with SR141716. Scale bars represent 10 ms (horizontal) and 50 pA (vertical). Data points and bar graphs represent mean ± SEM. Bonferroni-corrected or paired (vs. baseline) two-tailed t-test, **p < 0.01, ***p < 0.001, ****p < 0.0001.

We first hypothesized that CIE procedures induced a change in CB1 receptor function. We applied a CB1 receptor agonist (1 μM WIN 55,212-2) (Figure 2E, F), but found a similar decrease in EPSC amplitude between Air (44.72 ± 1.98% of baseline at 30-40 min, *n*=6, paired t test vs. baseline: t_5_ = 27.96, p < 0.0001) and CIE exposed mice (47.08 ± 9.03% of baseline at 30-40 min, *n* = 6, paired t test vs. baseline: t_5_ = 5.86, p < 0.01) (unpaired t test comparing Air and CIE mice: t10 = 0.25, p = 0.80).

Another possibility for the observed decrease in OFC transmission may involve a CIE-induced change in the expression of eCB-mediated plasticity (DePoy et al., 2013). Given the decreased OFC transmission induced by CIE, we hypothesized that eCB-mediated long-term depression (LTD) would be occluded in CIE exposed mice. eCB-LTD has been extensively studied and described using a high frequency stimulation (HFS) induction protocol. However, we needed to probe input-specific contributions, and conventional ChR2 shows a strong inactivation and desensitization to HFS (Lin, 2010). Thus, we opted to use a group 1 mGluR agonist, DHPG, to induce LTD (Kreitzer and Malenka, 2007, 2005; Wu et al., 2015; Yin et al., 2006). A 10-minute bath application of DHPG (50 μM) paired with postsynaptic depolarization to 0 mV, resulted in LTD of optically evoked EPSCs (Air: 54.60 ± 8.40% of baseline at 30-40 min, *n* = 6, paired t test vs. baseline: t_5_ = 5.41, p < 0.01; CIE: 46.68 ± 4.97% of baseline at 30-40 min, *n* = 5, paired t test vs. baseline: t_4_ = 10.74, p < 0.001) (Figure 2G). However, this DHPG-induced LTD was of similar magnitude between Air and CIE exposed mice (unpaired t test: t_9_ = 0.77, p = 0.46) (Figure 2H). Bath application of a CB1 receptor antagonist, SR141716 (1 μM) showed the observed LTD to be CB1 receptor dependent in both groups (Air: 103.2 ± 5.85% of baseline at 30-40 min, *n* = 4, paired t test vs. baseline: t_3_ = 0.54, p = 0.63; CIE: 108.3 ± 5.85% of baseline at 30-40 min, *n* = 6 paired t test vs. baseline: t_5_ = 0.96, p = 0.38) (Figure 2I, J). Thus, CB1 receptor dependent eCB-LTD at OFC-D1 SPNs synapses also remains intact after CIE exposure.

While CB1 receptor function and eCB-mediated plasticity at OFC-D1 SPN synapses was still intact in CIE mice, we wondered whether regulation of eCB signaling may be disrupted in alcohol dependence. Prior work in striatum has shown that activation of D1 receptors prevents the expression of eCB-LTD (Trusel et al., 2015; Wu et al., 2015). Additionally, positive-timing induction protocols in striatum require D1 receptor blockade for the expression of CB1 receptor dependent LTD (Shen et al., 2008). Based on these previous findings, we reasoned that D1 receptor activation may negatively regulate eCB-LTD. We found that activation of D1 receptors by bath application of SKF 81297 (3 μM) in Air controls blocked DHPG-LTD (96.32 ± 10.41% of baseline at 30-40 min, *n* = 6, paired t test vs. baseline: t_5_ = 0.35, p = 0.74) but had no effect in CIE exposed mice (56.12 ± 4.85% of baseline at 30-40 min, *n* = 7, paired t test vs. baseline: t_6_ = 9.06, p < 0.001). The LTD that CIE mice continued to express was still CB1 receptor dependent, as bath application of SR141716 (1 μM) blocked the DHPG-LTD (104.2 ± 11.55% baseline at 30-40 min, *n* = 5, paired t test vs. baseline: t4 = 0.37, p = 0.73) (Figure 2K, L).

This effect may not be specific to OFC-D1 SPNs and could also be present at any glutamatergic input onto D1 SPNs. However, evoked EPSCs by electrical stimulation in a subset of mice where OFC-DS transmission was optically assessed, showed a similar magnitude of DHPG-LTD in both Air and CIE exposed mice that was CB1 receptor dependent (Figure 2-figure supplement 2A-C). Further, DHPG-LTD of electrically evoked EPSCs was blocked by SKF 81297 in Air mice (Figure 2-figure supplement 2F, G). While activation of D1 receptors with bath application of DHPG did result in a short-term depression in CIE, electrically evoked EPSC amplitudes returned to baseline at 30-40 min (Figure 2-figure supplement 2F, G) in contrast to the long-lasting effect observed when optically evoking EPSCs (Figure 2K, L). Additionally, we found that this short-term depression was sensitive to a CB1 receptor antagonist, as we observed no short- or long-term changes in EPSC amplitude in the presence of SR141716 (1 μM) (Figure 2-figure supplement 2F, G). Thus, the disrupted D1 receptor regulation of long-term eCB plasticity appears to be at least somewhat selective to OFC-D1 SPNs, as we do not see similar changes when examining excitatory input generally. These results raise the hypothesis that D1 receptor signaling may be altered in CIE exposed mice.

### Ethanol dependence disrupts D1 receptor function

D1 receptors may negatively modulate eCB signaling as activation of D1 receptors increases spontaneous presynaptic neurotransmitter release (André et al., 2010), a decrease in eCB content (Patel, 2003), and prevents the expression of eCB-LTD (Wu et al., 2015). To determine whether CIE alters D1 receptor activity we used both an *in vitro* and *in vivo* assay of D1 receptor function. First, D1 receptor activation *in vitro* has been shown to increase excitability of D1 SPNs (Hernandez-Lopez et al., 1997; Planert et al., 2013; Surmeier et al., 2007). Similar to previous findings, we found that washing on D1 agonist, SKF 81297 (10 μM), enhanced D1 SPN action potential firing in response to current injections in the DS of Air controls (repeated-measures ANOVA: interaction (Current x SKF 81297): F_(10, 90)_ = 5.72, p < 0.0001; main effect of Current: F_(10, 90)_ = 22.46, p < 0.0001; main effect of SKF 81297: F_(1, 9)_ = 32.37, p = 0.0003) (Figure 3A). In contrast, bath application of SKF 81297 had no effect on D1 SPN excitability in CIE mice (repeated-measures ANOVA: no interaction (Current x SKF 81297): F_(10, 80)_ = 0.28, p = 0.98; main effect of Current: F_(10, 80)_ = 29.87, p < 0.0001 but no main effect of SKF 81297: F_(1, 8)_ = 0.09, p = 0.77) (Figure 3B). This suggests that D1 receptor signaling is disrupted in CIE mice and is in line with our previous data in which a D1 agonist failed to modulate DHPG-LTD in CIE mice (Figure 2K, L).

**Figure 3.**
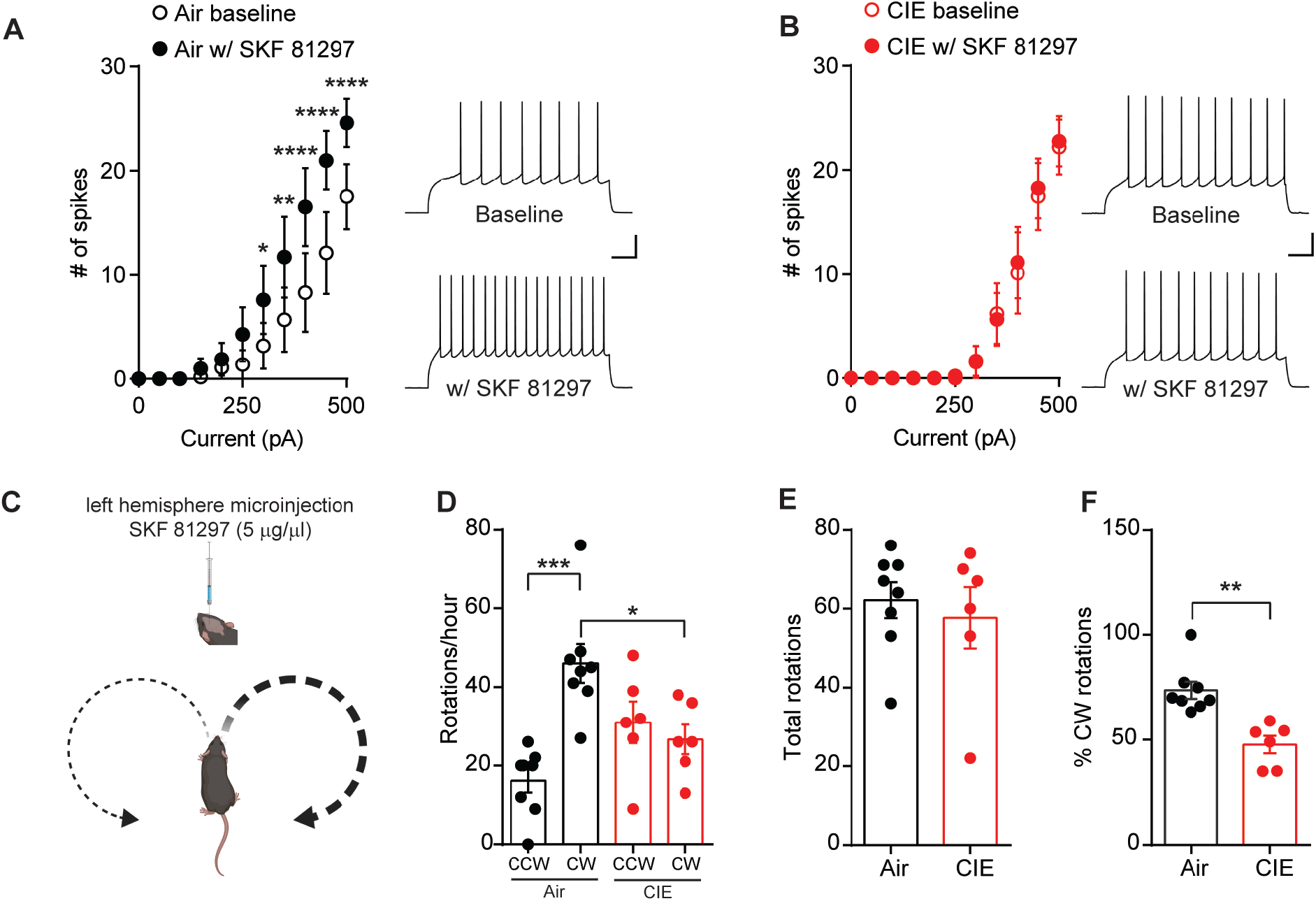
D1 receptor function is disrupted in CIE exposed mice. **(A)** Number of action potentials plotted against current injected under baseline conditions and in the presence of D1 agonist, SKF81297 (10 μM) (left) and sample traces (right) in D1 SPNs of Air (*n* = 10 cells, 3 mice) and **(B)** CIE (*n* = 9 cells, 4 mice). Scale bars represent 100 ms (horizontal) and 20 mV (vertical). **(C)** Schematic of unilateral microinjections (300 nL) of SKF 81297 (5 μg/μL) in the left hemisphere in the DS and predicted bias towards clockwise (CW) rotations. **(D)** Number of CW rotations and counterclockwise (CCW) rotations in Air (*n* = 8 mice) and CIE (*n* = 6 mice) (2 vapor cohorts) mice counted over an hour session. **(E)** Total rotations (CW+CCW) in Air and CIE mice. **(F)** Percentage of CW turns in Air and CIE mice. Data points and bar graphs represent mean ± SEM. Bonferroni-corrected post-hoc test of repeated-measures ANOVA or unpaired two-tailed t-test, *p < 0.05, **p < 0.01, ***p < 0.001, ****p < 0.0001.

To test whether alcohol dependence alters the function of DS D1 receptors *in vivo*, we relied on previous findings where an imbalance in striatal dopamine induced rotational behavior (Glick et al., 1976; Martín et al., 2008). We gave Air and CIE mice a unilateral microinjection of the D1 agonist, SKF 81297 (300 nL, 5 μg/μL), into the left hemisphere of the DS (Figure 3C, Figure 3-figure supplement 3) and we measured the subsequent number of clockwise (CW) and counterclockwise (CCW) rotations. The prediction is that an increase in unilateral dopamine signaling should normally lead to an increase in clockwise rotational behavior (Figure 3C). While Air mice showed the predicted increase in CW rotations, CIE mice showed reduced D1-agonist induced rotational behavior. A one-way ANOVA (F_(3, 24)_ = 9.31, p < 0.001) showed that Air controls had significantly more CW than CCW rotations (Bonferroni corrected, p < 0.001) and had more CW rotations compared to CIE mice (Bonferroni corrected, p < 0.05) (Figure 3D). Importantly, there was no difference in the total number of rotations between Air and CIE mice (Air: 62.13 ± 4.54, *n* = 8; CIE: 57.67 ± 7.76, *n* = 6; unpaired t test: t_12_ = 0.53, p = 0.61) (Figure 3E). However, the percentage of CW rotations out of total rotations in CIE mice was significantly lower compared to Air controls (Air: 73.72 ± 4.08, *n* = 8; CIE: 47.72 ± 4.21, *n* = 6; unpaired t test: t_12_ = 4.36, p < 0.001) (Figure 3F). The blunted D1 receptor agonist response in CIE mice could be due to alterations in D1 receptor expression; however, Western blot analysis showed no change in D1 receptor expression in the DS between CIE and Air mice (Figure 3-figure supplement 4). Together, these data provide evidence that CIE results in disruption of D1 receptor function downstream of D1 receptor activation.

### CIE enhances depolarization induced suppression of excitation of OFC transmission to D1 SPNs

Given the alcohol dependence-induced long-lasting disruption of D1 receptor function and the role of D1 receptors in the negative modulation of eCB signaling (André et al., 2010; Patel, 2003), we hypothesized that we may see an increase in eCB signaling in CIE mice. To test for CIE-induced changes in eCB signaling in OFC-D1 SPN transmission, we used depolarization-induced suppression of excitation (DSE), a short-term form of eCB plasticity (Diana and Marty, 2004). To induce DSE of optically evoked EPSCs from OFC terminals (Figure 4A), D1 SPNs were depolarized to +50 mV for 10 seconds which resulted in a short-term decrease of EPSC amplitudes in both Air (DSE Air: 25.78 ± 3.82, *n* = 8, 3 mice, paired t test vs. baseline: t_7_ = 6.75, p < 0.001) and CIE mice (DSE CIE: 42.10 ± 8.25, *n* = 9, 3 mice, paired t test vs. baseline: t_8_ = 15.31, p < 0.0001) that was CB1 receptor dependent (bath application of SR141716 (1 μM), DSE Air: 2.71 ± 3.09, *n* = 5, paired t test vs. baseline: t_4_ = 0.88, p = 0.43; DSE CIE: 1.77 ± 5.20, *n* = 6, paired t test vs. baseline: t_5_ = 0.34, p = 0.75) (Figure 4B, C). We found that CIE mice showed a larger magnitude of DSE than did Air controls (unpaired t test Air vs. CIE: t_15_ = 3.53, p < 0.01), indicating a CIE-induced increase in eCB signaling in OFC-D1 SPN transmission (Figure 4B, C). To again determine if this effect is selective for OFC-D1 input, we examined DSE in electrically evoked EPSCs (Figure 4D) and found DSE in both Air (DSE Air: 32.96 ± 2.50, *n* = 6, 3 mice, paired t test vs. baseline: t_5_ = 13.19, p < 0.0001) and CIE mice (DSE CIE: 36.84 ± 7.00, *n* = 6, 3 mice, paired t test vs. baseline: t_5_ = 5.27, p < 0.01), that was also sensitive to CB1 receptor blockade (DSE Air: 2.40 ± 4.88, *n* = 6, paired t test vs. baseline: t_5_ = 0.49, p = 0.64; DSE CIE: 3.97 ± 1.94, *n* = 5, paired t test vs. baseline: t_4_ = 2.05, p = 0.11). In contrast to optically evoked EPSCs at OFC-D1 SPNs synapses (Figure 4A-C), Air and CIE mice showed a similar magnitude of DSE when EPSCs were electrically evoked (unpaired t test Air vs. CIE: t_10_ = 0.52, p = 0.61) (Figure 4D-F). Further, the CIE-induced increase in DSE was cell-type selective as we observed no difference in DSE of D2 SPNs of Air and CIE mice with either optical stimulation of OFC terminals or local electrical stimulation (Figure 4-figure supplement 5). The CIE-induced increase in DSE was seen throughout the protracted withdrawal period (Figure 4-figure supplement 6). These results suggest that the increase in DSE of OFC transmission onto D1 SPNs following the induction of alcohol dependence are at least partially selective to the OFC input.

**Figure 4.**
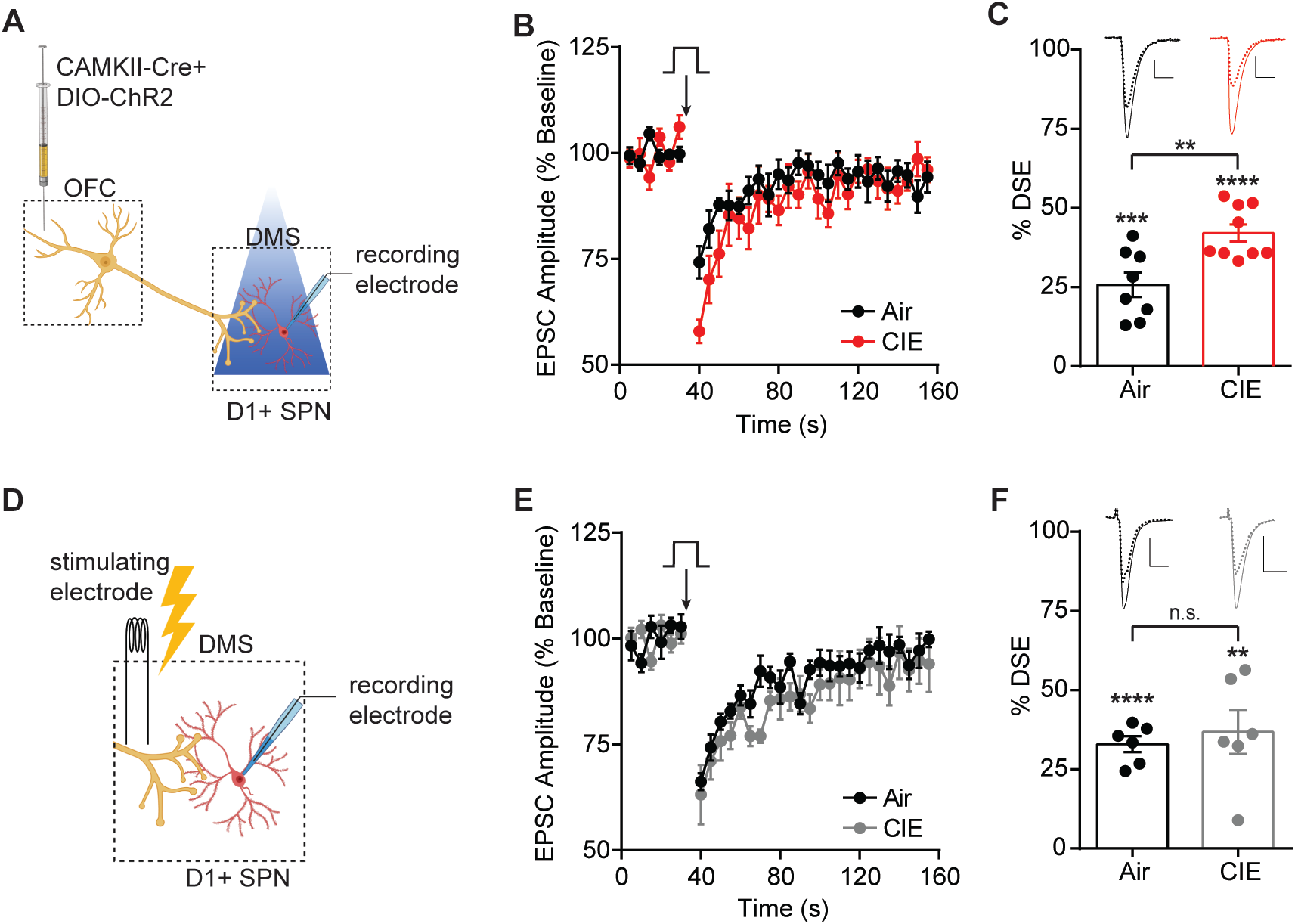
CIE induced enhancement of DSE at OFC terminals to D1 SPNs in the DS. **(A)** Schematic of viral injections in OFC and optical stimulation of OFC terminals during whole-cell recordings of D1 SPNs in the DS. **(B)** Depolarization induced suppression of excitation (DSE) of OFC inputs to D1 SPNs in Air (*n* = 8 cells, 3 mice) and CIE (*n* = 9 cells, 3 mice). **(C)** Bar graphs representing the percentage change from baseline immediately after depolarization. **(D)** Schematic of local electrical stimulation during whole-cell recordings of D1 SPNs in the DS. **(E)** DSE of excitatory inputs to D1 SPNs using electrical stimulation in Air (*n* = 6 cells, 2 mice) and CIE (*n* = 6 cells, 3 mice). **(F)** Bar graphs representing the percentage change from baseline immediately after depolarization. Scale bars represent 10 ms (horizontal) and 50 pA (vertical). Data points and bar graphs represent mean ± SEM. Scale bars represent 10 ms (horizontal) and 50 pA (vertical). Unpaired (Air vs. CIE) or paired (vs. baseline) two-tailed t-test, **p < 0.01, ***p < 0.001, ****p < 0.0001.

### CB1 receptor antagonist restores OFC transmission and goal-directed control

The above data suggests an enhancement of eCB signaling at OFC-D1 SPN synapses in CIE mice. Given this, we reasoned that CB1 receptor antagonism may alter the aberrant *in vivo* activity seen in OFC-DS terminals and restore sensitivity to outcome devaluation. However, the prevalence of CB1 receptors in cortico-basal ganglia circuits makes indirect actions on circuit activity also likely, which could obscure or counter effects at OFC-DS terminals. To examine this, we gave systemic injections of Vehicle or CB1 antagonist, SR141716 (3 mg/kg), prior to operant sessions and measured subsequent calcium activity in OFC-DS terminals during behavior (Figure 5A). Blocking CB1 receptors does not interfere with the ability to lever press (Hilario et al., 2007), and we also found that systemic administration of SR141716 had no obvious effect on gross lever press-related behavior (Figure 5-figure supplement 7).

**Figure 5.**
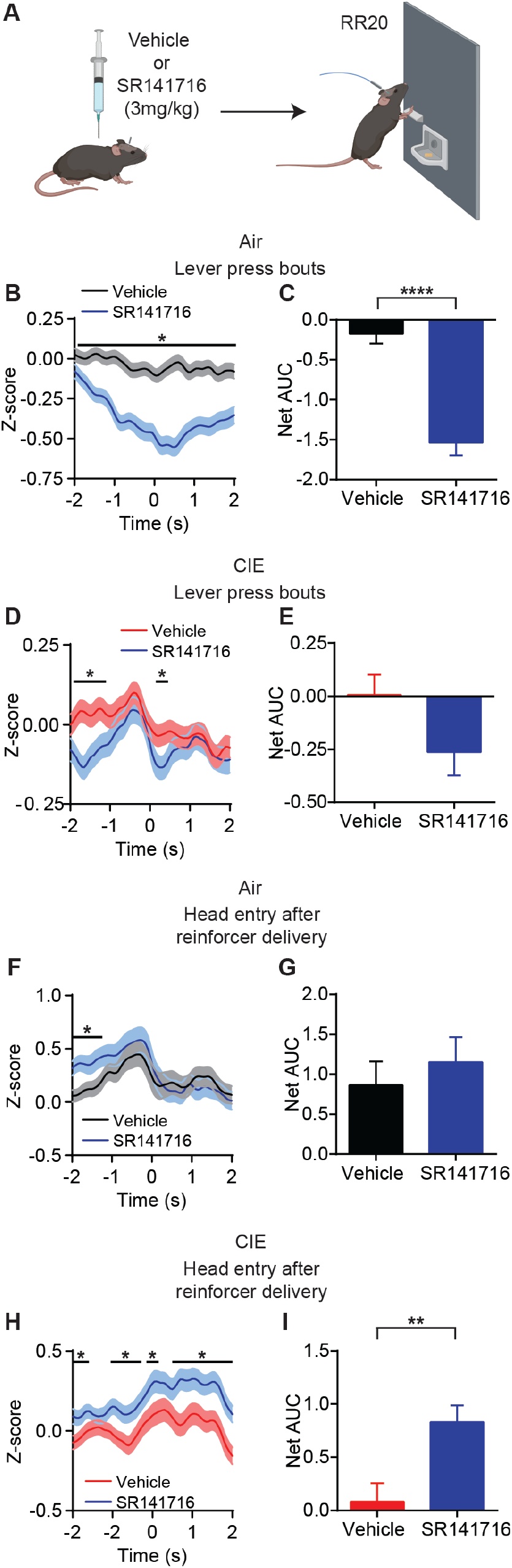
CB1 receptor antagonist restores OFC terminal activity for outcome encoding. **(A)** Schematic that depicts injections with vehicle or SR141716 followed by an operant session under a random ratio 20 (RR20) schedule of reinforcement in Air (*n* = 9 mice, 3 vapor cohorts) and CIE (*n* = 11 mice, 3 vapor cohorts). **(B)** Z-score of ΔF/F GCaMP6s trace recorded in OFC terminals during lever press bouts in mice injected with vehicle or SR141716 and **(C)** net area under the curve (AUC) for each signal in Air and **(D)**, **(E)** CIE mice. **(F)** Z-score of ΔF/F GCaMP6s trace recorded in OFC terminals during the first head entry after reinforcer delivery in mice injected with vehicle or SR141716 and **(G)** net area under the curve (AUC) for each signal in Air and **(H)**, **(I)** CIE mice. Data points and bar graphs represent mean ± SEM. Two-sided FDR-corrected permutation test or unpaired two-tailed t-test, *p < 0.05, **p < 0.01, ****p < 0.0001

We first looked at terminal activity during lever press bouts, where we previously saw slightly enhanced activity in CIE mice (Figure 1I, J). In Air control mice, systemically blocking CB1 receptors decreased OFC terminal activity during lever press bouts (FDR-corrected two-sided permutation test, Vehicle: *n* = 462; SR141716: *n* = 359) (Figure 5B, C). This was also reflected in the observed significant decrease in Net AUC (Vehicle: −0.21 ± 0.12, *n* = 462; SR141716: −1.42 ± 0.16, *n* = 359; unpaired t test: t_819_ = 6.27, p < 0.0001) (Figure 5C). Similarly, CIE mice also showed a decrease in OFC terminal activity prior to lever press bout onset with CB1 receptor blockade (FDR-corrected two-sided permutation test, Vehicle: *n* = 536; SR141716: *n* = 385) and but no significant decrease in Net AUC (Vehicle: 0.04 ± 0.09, *n* = 536; SR141716: −0.17 ± 0.14, *n* = 385; unpaired t test: t_919_ = 1.50, p = 0.14) (Figure 5D, E). As blocking CB1 receptors at OFC-DS terminals should prevent any eCB-related decrease in calcium activity (Kreitzer and Regehr, 2001), this suggests the reduced lever-press related OFC-DS terminal activity is due to eCB actions on broader circuit activity.

However, eCBs are thought to be released at glutamatergic synapses in DS in an activity-dependent manner (Adermark et al., 2009; Adermark and Lovinger, 2009) and we saw larger increases in OFC-DS terminal activity during periods of associated with outcome retrieval (Figure 1L, M). Hence, we also examined effects of blocking CB1 receptors on the reduced outcome retrieval activity observed in CIE mice (Figure 1P, Q). Systemic administration of the CB1 antagonist in Air controls resulted in a brief, slight increase OFC terminal activity prior to the first head entry after reinforcer delivery (FDR-corrected two-sided permutation test, Vehicle: *n* = 177; SR141716: *n* = 162) (Figure 5F), but did not induce a difference in Net AUC (Vehicle: 1.05 ± 0.28, *n* = 177; SR141716: 1.15 ± 0.32, *n* = 162; unpaired t test: t_337_ = 0.24, p = 0.81) (Figure 5G). In contrast, systemic administration of the CB1 antagonist in CIE mice produced a significant and sustained increase in OFC terminal activity during this period of outcome retrieval (FDR-corrected two-sided permutation test, Vehicle: *n* = 275; SR141716: *n* = 219) (Figure 5H) and a significant increase in net AUC (Vehicle: 0.09 ± 0.18, *n* = 275; SR141716: 0.83 ± 0.16, *n* = 219; unpaired t test: t_492_ = 3.10, p < 0.01) (Figure 5I). A similar pattern was observed when we examined effects of the CB1 antagonist on average animal session data (Figure 5-figure supplement 8). Thus, it appears that systemic administration of a CB1 antagonist produces different effects on OFC-DS terminal activity dependent on the behavior. Notably, we see a large CB1 receptor antagonist-induced increase in OFC-DS terminal activity during periods associated with outcome retrieval in CIE mice, in which OFC-DS terminal is normally blunted and endocannabinoid signaling is enhanced.

Given this, we hypothesized that blocking eCB signaling in DS would restore adaptive outcome devaluation in alcohol dependent mice. Two separate cohort replicates of mice were implanted with bilateral cannula targeting DS (Figure 6-figure supplement 9) and then exposed to Air or CIE followed by lever press training for a food outcome (Figure 6A, B) as previously described (Figure 1). Air and CIE mice pressed the lever at similar levels (repeated-measures ANOVA: main effect of Training day: F_(8, 240)_ = 161.5, p < 0.0001; no main effect of CIE exposure or interaction: Fs < 0.13) (Figure 6C) and earned a similar number of reinforcers across training (repeated-measures ANOVA: main effect of Training day: F_(8, 240)_ = 71.45, p < 0.0001; no main effect of CIE exposure or interaction: Fs < 0.18) (Figure 6D). As described above, lever presses were organized into bouts (Figure 1). Similar to the previous cohorts, we found that lever press bouts were differentially executed in Air and CIE mice in that CIE mice had slightly less presses per bout, shorter inter-press intervals and consequently, shorter bout durations (Figure 6-figure supplement 9).

**Figure 6.**
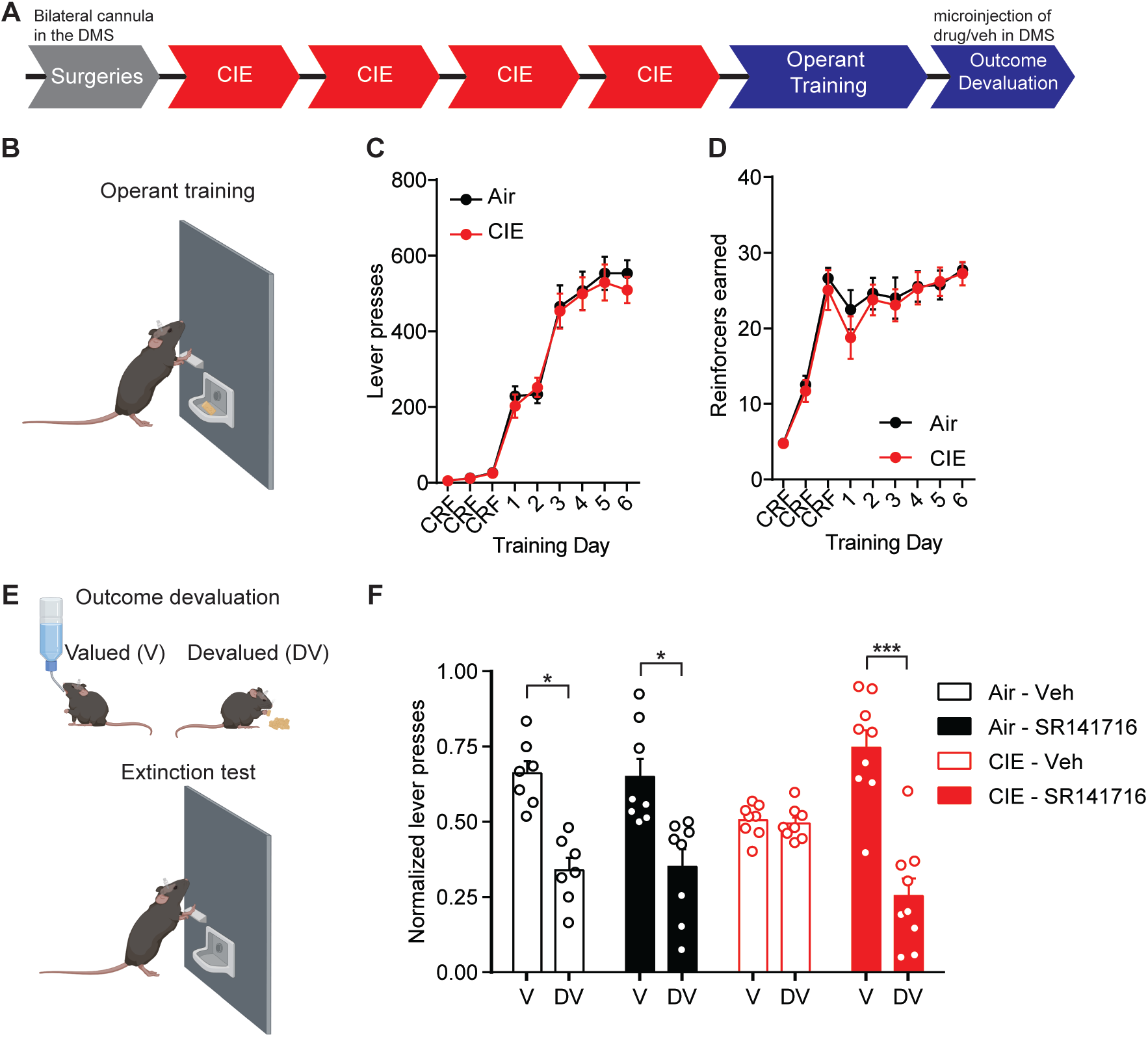
Blockade of CB1 receptors in the DS restores value-based decision-making in CIE mice. **(A)** Experimental timeline that includes surgeries followed by 4 cycles of CIE exposure, operant training and outcome devaluation **(B)** Schematic of mouse lever press training in which Air (Vehicle: *n* = 7 mice, SR141716: *n* = 8 mice, 3 vapor cohorts) and CIE mice (Vehicle: *n* = 8 mice, SR141716: *n* = 9 mice, 3 vapor cohorts) were trained under a random ratio schedule of reinforcement for a food reinforcer. **(C)** Lever presses during operant training in Air and CIE mice. **(D)** Reinforcers earned during operant training in Air and CIE mice. **(E)** Schematic of outcome devaluation procedure that includes one-hour access to sucrose (valued) or the reinforcer earned during training (devalued) followed by a 5 min extinction test. **(F)** Normalized lever presses showing the distribution of lever pressing between the valued and devalued day in Air and CIE mice that received microinjections (300 nL) of vehicle or SR141716 (2 μM). Data points and bar graphs represent mean ± SEM. Bonferroni-corrected post-hoc test of repeated-measures ANOVA, *p < 0.05, ***p < 0.001.

We then examined the sensitivity of lever pressing to changes in expected outcome value using outcome devaluation procedures (Figure 6E). Just prior to outcome devaluation procedures, mice were injected with either vehicle or CB1 antagonist, SR141716, directly in the DS. We saw that in Air mice, intra-DS SR141716 had no effect on outcome devaluation responding. Once again, CIE exposed mice treated with vehicle did not show sensitivity to outcome devaluation. However, CIE mice that were given an injection of SR141716 in the DS significantly decreased lever press responding on the devalued day relative to the value day (Figure 6F). A repeated measures ANOVA, showed a significant interaction (Training day x Treatment: F_(3, 28)_ = 4.41, p = 0.01) and main effect of Devaluation day (F_(1, 28)_ = 33.28, p < 0.0001) but no main effect of Treatment (F < 0.1). Thus, locally blocking CB1 receptors in the DS of CIE mice was sufficient to restore value-based decision-making.

## Discussion

Often differing symptoms in psychiatric conditions produce opposing changes in the activity of a brain area. As there is increasing potential for cortical activity modulation to be used in the treatment of psychiatric disorders, we need a greater understanding of how such cortical modulation will affect downstream circuit engagement and information representation. Here we find that alcohol dependence enhances activity associated with actions in OFC terminals in striatum, while also reducing activity associated with outcome retrieval. We identify one mechanism potentially responsible for the reduced OFC-DS activity specific to epochs of outcome retrieval: a disruption in D1 receptor function in SPNs and an enhancement of eCB signaling that reduces OFC-D1 SPN transmission in the DS (Figure 7). Restoration of OFC-DS transmission through CB1 receptor blockade restored use of reward value to control decision-making. Thus, alcohol dependence induces long-lasting, computational-specific changes to cortical transmission in part through cell-type, synapse specific changes in post-synaptic modulation of this cortical terminal release.

**Figure 7.**
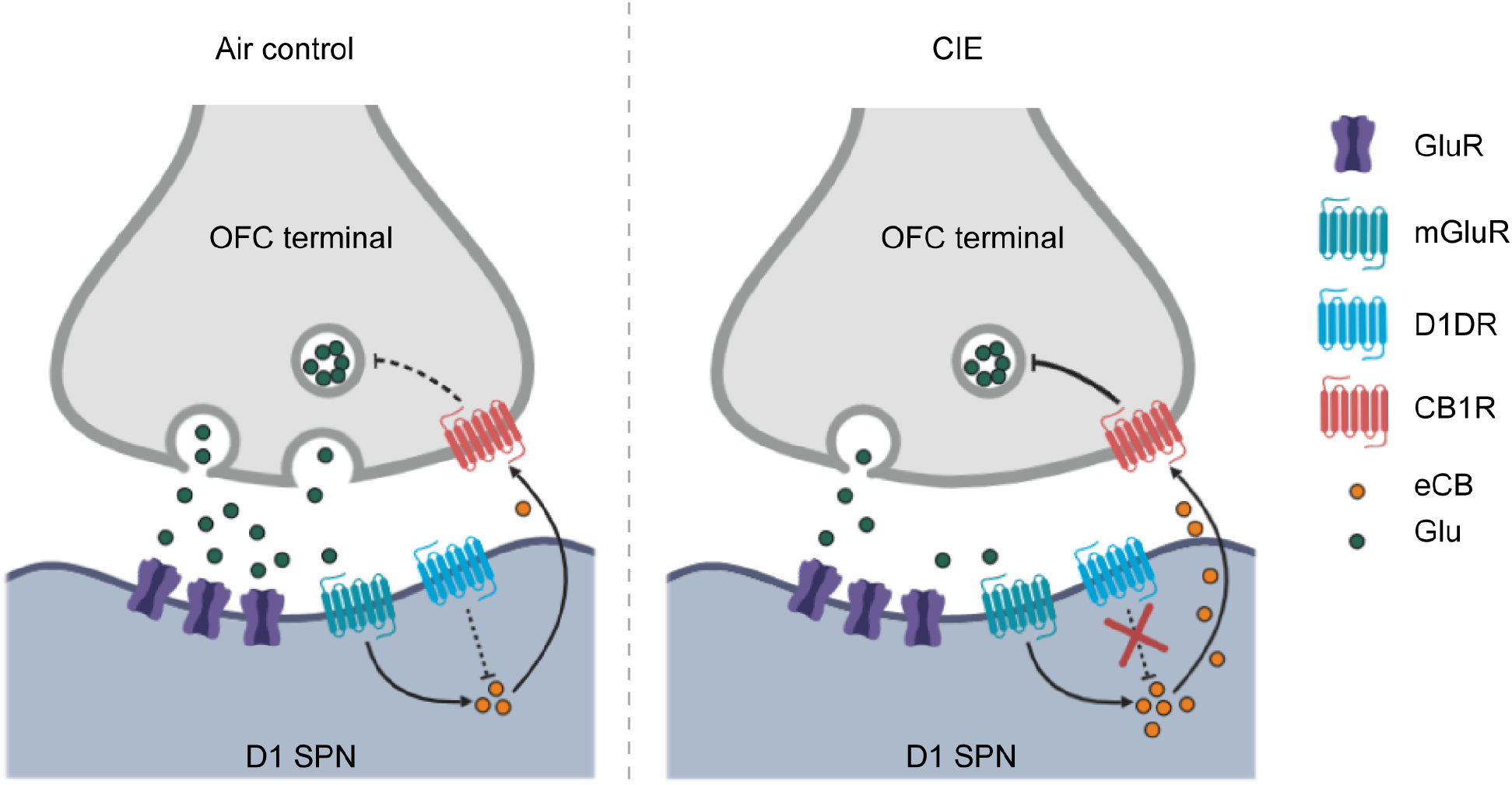
Summary of hypothesized alterations in D1 receptor function and endocannabinoid signaling in D1 SPNs of CIE exposed mice. In Air controls (left), D1 receptor activity regulates endocannabinoid production and/or release. In CIE mice (right), the loss of D1 receptor function leads to unregulated endocannabinoid signaling and the long-term reduction in glutamate release from OFC inputs.

We observed OFC-DS terminal activity profiles reminiscent of common OFC observations in humans: OCD-related hyperactivity (Milad and Rauch, 2012; Pauls et al., 2014; Robbins et al., 2019), the hallmark representation of reward evaluation (Fellows, 2007; Wallis, 2007), and dependence-induced hypoactivity (Catafau et al., 1999; Lüscher et al., 2020; Schoenbaum et al., 2016, 2006; Volkow et al., 1997; Volkow and Fowler, 1994). Alcohol dependent mice showed slightly increased OFC terminal activity during lever press behavior (Figure 1), similar to the increased OFC-DS activity reported in compulsive phenotypes (Pascoli et al., 2018). While we did not directly measure markers of compulsivity in the present studies, we have previously reported that CIE procedures can increase response rates for both food reinforcers (Renteria et al., 2018) and alcohol self-administration (Renteria et al., 2020). This raises the hypothesis that OFC-DS transmission has a role in alcohol-induced compulsive seeking. However, while this increased OFC terminal activity may drive the increase in response rate in CIE mice, it may not influence reward evaluation processes. We previously observed no correlation between response rate and the sensitivity to outcome devaluation following alcohol dependence (Renteria et al., 2018). Hence action control and the evaluation of outcome value may be differentially affected by alcohol dependence. That we also found reduced OFC-DS terminal activity in CIE mice during periods associated with outcome retrieval shows how population output activity may be differentially modulated depending on the behavior measured. The observations that alcohol dependence significantly decreased reward-related OFC-DS terminal activity, but increased action-related activity, raises one hypothesis that alcohol-dependence differentially alters afferent activity into OFC thereby altering the capability of OFC-DS projections to send action and outcome information into basal ganglia circuits. Further, the restoration of OFC terminal activity with a CB1 antagonist during outcome retrieval (Figure 5) and rescue of goal-directed control when injected directly in DS (Figure 6), suggests there also is additional local synaptic modulation of OFC-DS terminal activity during behavior.

Our data suggests that dependence induces an increase in eCB signaling at OFC-D1 SPN synapses. While multiple aspects of eCB downstream signaling and plasticity were still intact, including CB1 receptor function and the ability to induce an eCB dependent LTD (Figure 2), upstream regulation of eCB signaling by D1 receptors was not. Using *ex vivo* brain slice preparations from CIE mice, we observed no effect of the D1 agonist SKF 81297 on DHPG-LTD or the excitability of D1 SPNs as previously reported (Hernandez-Lopez et al., 1997; Planert et al., 2013; Surmeier et al., 2007) and observed in our controls. *In vivo* unilateral microinjections of SKF 81297 resulted in rotation behavior that was significantly attenuated in CIE mice and there was no change in D1 receptor expression (Figure 3-figure supplement 4). These results suggest that CIE disrupts the signaling cascade downstream of D1 receptor activation. Indeed, acute ethanol has been shown to modulate downstream targets of D1 receptor activation (Ron and Barak, 2016; Ron and Messing, 2013), including cellular inhibition of specific protein kinase C (PKC) isoforms that constitutively phosphorylate D1 receptors to dampen D1 signaling (Rex et al., 2008). Withdrawal was found to decrease PKC isoforms in the BLA (Christian et al., 2012), although whether similar mechanisms occur in dorsal striatum is not known. As our measures of D1 receptor function were done 3-21 days in withdrawal from chronic ethanol exposure, this evidence raises one hypothesis that the present disruption in D1 receptor function observed may reflect neuroadaptive compensatory changes in PKC activity. Others have shown, in postmortem brain sections taken from human alcoholics, a down-regulation of D1 receptor binding sites in the striatum (Hirth et al., 2016). In addition, rats exposed to CIE had an increase of dopamine release in the nucleus accumbens shell, but showed a blunted response to D1 receptor activation 21 days in withdrawal (Hirth et al., 2016). While plenty of data suggest that chronic alcohol dependence alters dopamine systems, the changes likely depend on the synapse, cell-type, and circuit.

The loss of D1 receptor function in SPNs of CIE exposed mice may be directly related to the enhancement of eCB signaling (Figure 4 and Figure 7). Previous works have shown dopamine and eCB signaling interact in which D1 receptor activation may reduce anandamide (AEA) signaling (André et al., 2010; Patel, 2003) (but see (Giuffrida et al., 1999)), although, at what point D1 signaling could interfere with AEA synthesis or release is not clear. However, DSE has widely been shown to be mediated by 2-arachidonoyl glycerol (2-AG) and a CIE-induced increase in 2-AG may contribute to the present effects. Others have shown that habitual alcohol seeking may be mediated by an increase in 2-AG, as an inhibitor, DO34, was found to reduce habitual responding (Gianessi et al., 2020). Thus, future work examining the contribution of different eCBs is warranted.

Conversely, the eCB system has been extensively studied in the modulation of dopamine transmission (Covey et al., 2017), further underscoring the complex interactions of these signaling systems. For example, as eCBs can reduce dopamine release in nucleus accumbens through reducing cortical drive (Mateo et al., 2017), there may also be a CIE-induced decrease in endogenous D1R activation at OFC-D1 SPN synapses. Given the large number of synaptic and molecular targets that chronic ethanol exposure can modulate (Abrahao et al., 2017; Lovinger and Roberto, 2013; Roberto and Varodayan, 2017), the synapse and cell-type specific alterations in orbitostriatal transmission that we have observed are likely the result of ethanol-induced changes in many neuromodulatory and synaptic signaling systems.

The CIE-induced alterations in eCB-mediated signaling and plasticity presented here were found to be at least in part, specific to the OFC input. Local electrical stimulation (Figure 2-figure supplement 2) likely includes OFC inputs in addition to excitatory inputs from other cortical and limbic regions as well as thalamic nuclei (Wall et al., 2013). It is possible that other excitatory inputs to D1 SPNs are differentially modulated by alcohol dependence such that changes at individual inputs may be masked with electrical stimulation. However, enhanced eCB signaling is not expected to apply to thalamostriatal inputs to D1 SPNs, as they lack CB1 receptors at presynaptic terminals (Wu et al., 2015). Evidence for a CIE-induced change in other excitatory inputs to D1 SPNs may be supported by the DHPG-induced short-term depression observed with electrical stimulation in the presence of a D1 agonist (Figure 2-figure supplement 2). As the observed short-term depression was found to be sensitive to a CB1 receptor antagonist, it likely involves a change in eCB signaling, although the mechanism may differ from that observed with long-term depression (Castillo et al., 2012; Heifets and Castillo, 2009; Kano et al., 2009; Lovinger, 2008).

Interestingly, we found the effects of CB1 antagonist on OFC terminal activity to be dependent on the specific component of decision-making represented. Systemic injection of CB1 receptor antagonist was found to decrease OFC terminal activity in DS during initiation of lever press bouts in both Air and CIE mice (Figure 5), suggesting the effect the antagonist exerted was not through activation of CB1 receptors on OFC terminals. One explanation could be a disinhibition of inhibitory signaling onto OFC cell bodies or terminals in DS. CB1 receptors are expressed on GABAergic terminals in cortex which, when blocked with a CB1 antagonist, would allow for inhibition of cortical projection neurons. Additionally, in the striatum CB1 receptors are expressed not only on cortical terminals, but also on SPNs and GABAergic interneurons (Shu-Jung Hu and Mackie, 2015) which may provide local regulation of excitatory transmission. In contrast, blocking CB1 receptors had little effect on OFC-DS transmission during outcome retrieval epochs in Air controls but significantly increased activity in CIE mice. These observations suggest that differences in instrumental responding and outcome encoding may result from differentially engaged local regulation of OFC-DS transmission. As prior work showed a dependency of outcome devaluation on the activity of OFC-DS terminals (Gremel et al. 2016), it is important to note that our current findings suggest that OFC may pass into striatum information pertaining to more than one type of computation employed during outcome devaluation.

The present mechanistic findings, which are not only synapse, cell-type, and projection specific, but likely also dependent on input engagement or computation performed, demonstrate the complexity of how circuits represent information. A circuit informational approach (Lovinger & Gremel, 2020), aimed at identifying the strength and pattern of incoming transmission, recruited plasticity mechanisms, and capabilities of local circuit modulation to affect transmission, increasingly appears necessary in order to understand disease-induced disruptions to behavior. Here we further our understanding of how OFC-DS circuitry regulates decision-making, building off of previous works (Gremel & Costa, 2013; Gremel et al., 2016) to show this pathway carries both action and reward information. Importantly, we now show that action and reward information can recruit different downstream mechanisms depending on the pattern of activation. However, much work remains to be done to understand the neuroadaptations induced by CIE that result in aberrant decision-making. For example, what does the reduction in transmission from one cortical area mean for SPN output in the DS? Results in the present study focus on the effects of CIE on OFC-D1 SPN transmission in the DS but the loss of value-based decision-making likely also involves changes to habitual processes for which the neurocircuitry is less understood. Work performed in non-human primates suggests chronic heavy alcohol consumption increases excitatory drive and decreases local inhibition within the putamen or dorsolateral striatum (DLS), an area supporting habitual action control (Cuzon Carlson et al., 2011). Thus, it may be that alcohol dependence results in a decreased ability of associative cortical input to control DS output as well as disinhibition of DLS. Elucidating the effects of alcohol dependence on corticostriatal plasticity will further our understanding of the dependence-induced disruptions to decision-making processes that contribute to continual drug seeking and taking behaviors.

## Methods

### Mice

B6.Cg-Tg(Drd1a-tdTomato)6Calak/J (Shuen et al., 2008) and B6.FVB(Cg)-Tg(Drd1-cre)EY266Gsat/Mmucd mice were used for electrophysiological recordings and C57BL/6J mice were used for behavioral experiments and fiber photometry. All mouse lines were obtained from Jackson Laboratory and transgenic mouse lines were bred with C57Bl/6J mice (Jackson Laboratory) for one generation, in-house. Adult (> 8 weeks) male and female mice were housed in groups of 2-4, with mouse chow (Purina 5015) and water ad libitum and were kept on a 14-hour light/10-hour dark cycle. Behavioral and physiology experiments were conducted during the light phase of the light cycle. All experiments were approved by the Institutional Animal Care and Use Committees of the University of California San Diego and experiments were conducted according to NIH guidelines.

### Viral injections

Mice were anesthetized with isoflurane and were given stereotaxically guided bilateral injections into the OFC (coordinates from Bregma in mm: anterior [A], 2.70; medial [M], ±1.65; ventral [V]: 2.65). D1-tdTomato mice, used for patch clamp recordings, were co-injected with 100 nL AAV5-CamKIIa-GFP-Cre and 100 nL AAV5-Ef1a-DIO-ChR2-eYFP in the OFC. D1-Cre mice received additional bilateral injections of AAV5-Flp-tdTomato in the DS (coordinates from Bregma in mm: anterior [A], 0.5; medial [M], ±1.5; ventral [V]: 3.25). The coordinates target the medial DS from the perspective of the dorsal-ventral axis, and more medial DS in respect to the medial versus lateral striatal distinctions. For fiber photometry, C57BL/6J mice received unilateral injections with 500 nL AAV-hSyn-axon-GCaMP6s in the OFC. Viral spread and expression in OFC terminals were assessed by imaging the extent of fluorescence in brain slices (Olympus MVX10).

### Chronic intermittent ethanol exposure and repeated withdrawal

Mice were exposed to 4 rounds of ethanol vapor or air and repeated withdrawal (Becker, 1994; Becker and Lopez, 2004; Griffin 3rd et al., 2009; Lopez and Becker, 2005). Each round consisted of 16 h of vapor exposure followed by an 8 h withdrawal, repeated for 4 consecutive days. Ethanol was volatilized by bubbling air through a flask containing 95% ethanol at a rate of 2–3 L/min. The resulting ethanol vapor was combined with a separate air stream to give a total flow rate of approximately 10 L/min, which was delivered to the mice housed in Plexiglas chambers (Plas Labs Inc). Blood ethanol concentrations (BEC) were collected at the end of each round from sentinel mice (mean BEC = 34.7 ± 2.0 mM). No pyrazole or loading ethanol injections were given prior to placement in vapor chambers (Renteria et al., 2018).

### Operant behavior

Training was conducted as previously described (Gremel et al., 2016; Gremel and Costa, 2013; Renteria et al., 2018). Two days prior to training, mice were food restricted and maintained at ~ 90% of their baseline body weight throughout training and testing. Mice were placed in sound attenuating operant boxes (Med-Associates) and were trained to press a single lever (left or right) for a food reinforcer (chow pellet). Mice were first trained to retrieve the outcome, in the absence of levers, under a random time schedule (RT60) in which the outcome was delivered on average every 60 seconds. Mice were then trained on a continuous reinforcement schedule (CRF) in each context in which each lever press produced a single outcome, and the maximum number of reinforcers earned in 3 daily sessions being 5, 15, and 30, respectively. Following CRF training, mice were trained under a random ratio (RR) schedule of reinforcement. Mice received two days of training in RR10 (on average the 10^th^ lever press produces the outcome), followed by four days under RR20. Sessions ended after 30 reinforcers were earned or after 60 min had elapsed. Lever press bouts were analyzed using custom scripts on Matlab. Briefly, we found the inter press intervals for all lever presses within a session and calculated the mean. Bouts were separated by inter press intervals that were greater than the mean inter press interval. Only bouts of at least 3 lever presses were included for analysis.

Devaluation testing through sensory-specific satiation was conducted across two days: a valued day and a devalued day. Mice were allowed to prefeed ad libitum for 1 hour on the food pellet outcome previously earned by lever pressing (devalued day) or a 20% sucrose solution to control for satiety (valued day). Each day immediately following prefeeding, mice underwent a 5-minute extinction test in which the number of lever presses made were recorded, but no outcome was delivered. Lever presses on the valued or devalued day were normalized to total lever presses (valued + devalued) across devaluation testing to account for differences in response rates across individual groups. Mice with optical fiber implants for OFC terminal recordings were retrained after outcome devaluation using RR20 schedule of reinforcement and showed response rates during retraining similar to responding prior to outcome devaluation (Figure 5-figure supplement 6).

### Fiber photometry

Mice were injected with AAV-hSynapsin-axon-GCaMP6s in the OFC and fibers were implanted in the DS (coordinates from Bregma in mm: anterior [A], 0.5; medial [M], −1.5; ventral [V]: 3.25). Fibers were attached for the last 5 days of operant training, beginning on the 2^nd^ day of random ratio training. A blue LED (470 nm, Thorlabs) was used for excitation of OFC terminals in the DS. Fluorescence emissions were collected with a bifurcated fiber (Thorlabs) which allowed for simultaneous recording of two mice. The dual fiber core was focused through a 10x objective (Olympus) onto a CMOS camera (FLIR Systems). Fluorescence intensity and analog signals for lever press, head entries and reinforcer delivery were acquired simultaneously, thresholded, and timestamped for later analyses using Bonsai software(Lopes et al., 2015). Photometry and behavioral data were imported to Matlab (Mathworks Inc., Natick, MA, USA) and were analyzed using custom scripts. To account for photobleaching and signal decay across a session, we fit a double exponential line on the raw calcium fluorescence signal and normalized the signal to this fit. We then estimated baseline fluorescence by calculating the running 10^th^ percentile of the fluorescence signal using a 15 second sliding window. Only sessions in which more than 97.5% of the total session’s fluorescence signal exceeded a 1% change from baseline fluorescence were included (Markowitz et al., 2018). For each session, “trials” were composed of peri-event signals from −5 to 5 seconds around event onset. In each “trial”, 50 ms bins were z-scored to the session’s mean pre-event baseline (−5 to −2 seconds). We analyzed this photometry data two ways. First, these z-scored traces were combined across all mice within a group. This was done to conserve the variance seen within a subject. Second, we averaged these z-scored traces for a given animal session, and then averaged these traces across mice within a group. This examines between-mouse variability but does not preserve within-subject variability. Outliers, defined as peak amplitude Z-scores more than three scaled median absolute deviations (above and below) from the median, were removed from group data.

### Brain slice preparation

Mice were at least 12 weeks of age at the time of slice preparation. Coronal slices (250 μm thick) containing the DS were prepared using a Pelco easiSlicer (Ted Pella Inc., Redding, CA). Mice were anesthetized by inhalation of isoflurane and brains were rapidly removed and placed in 4°C oxygenated ACSF containing the following (in mM): 210 sucrose, 26.2 NaHCO_3_, 1 NaH_2_PO_4_, 2.5 KCl, 11 dextrose, bubbled with 95% O_2_/ 5% CO_2_. Slices were transferred to an ACSF solution for incubation containing the following (in mM): 120 NaCl, 25 NaHCO_3_, 1.23 NaH_2_PO_4_, 3.3 KCl, 2.4 MgCl_2_, 1.8 CaCl_2_, 10 dextrose. Slices were continuously bubbled with 95% O_2_/ 5% CO_2_ at pH 7.4, 32°C, and were maintained in this solution for at least 60 minutes prior to recording.

### Patch clamp electrophysiology

Whole-cell patch clamp recordings were made in identified SPNs in the medial DS. Cells were identified using an Olympus BX51WI microscope mounted on a vibration isolation table. Prior to patching onto a cell, the presence of td-Tomato expression was used to verify cell-type (D1+ or D1-) as well as eYFP expression for terminal expression of ChR2. eYFP expression was never observed in SPN cell bodies. Recordings were made in ACSF containing (in mM): 120 NaCl, 25 NaHCO_3_, 1.23 NaH_2_PO_4_, 3.3 KCl, 0.9 MgCl_2_, 2.0 CaCl_2_, and 10 dextrose, bubbled with 95% O_2_/ 5% CO_2_. ACSF was continuously perfused at a rate of 2.0 mL/min and maintained at a temperature of 32°C. Picrotoxin (50μM) was included in the recording ACSF to block GABAA receptor-mediated synaptic currents. Recording electrodes (thin-wall glass, WPI Instruments) were made using a PC-10 puller (Narishige International, Amityville, NY) to yield resistances of 3-6 MΩ. For current clamp experiments, electrodes were filled with (in mM): 135 KMeSO_4_, 12 NaCl, 0.5 EGTA, 10 HEPES, 2 Mg-ATP, 0.3 Tris-GTP, 260-270 mOsm (pH 7.3). For voltage clamp experiments, electrodes were filled with (in nM): 120 CsMeSO_4_, 15 CsCl, 8 NaCl, 10 HEPES, 0.2 EGTA, 10 TEA-Cl, 4 Mg-ATP, 0.3 Na-GTP, 0.1 spermine, and 5 QX-314-Cl. Access resistance was monitored throughout the experiments. Cells in which access resistance varied more than 20% were not included in the analysis.

Glutamatergic afferents were stimulated either electrically or optically. For electrical stimulation, a stainless steel bipolar stimulating electrode (FHC, Inc.) was placed dorsal to the recording electrode, about 150-300μm from the cell body. Optical stimulation was done using 470 nm blue light (4 ms pulse width) delivered via field illumination using a high-power LED (LED4D067, Thor Labs). Light intensity was adjusted to produce optically evoked excitatory postsynaptic currents (EPSCs) with a magnitude of 100–300 pA. Recordings were made using a MultiClamp 700B amplifier (Molecular Devices, Union City, CA), filtered at 2 kHz, digitized at 10 kHz with Instrutech ITC-18 (HEKA Instruments, Bellmore, NY), and displayed and saved using AxographX (Axograph, Sydney, Australia). For long term depression (LTD) experiments, EPSCs were evoked at 0.05 Hz for a 10 min baseline, followed by a 10 min bath application of (S)-3,5-Dihydroxy-phenylglycine (S-DHPG) (50uM) paired with depolarization to 0 mV. After S-DHPG application, EPSCs were monitored for 20 minutes at a rate of 0.05 Hz. Data were combined in 2 min bins and EPSC amplitudes were normalized to the 10 min baseline period. The magnitude of LTD was calculated by averaging normalized EPSC values from 30 to 40 min. For current-clamp recordings, a fixed current was injected for 800 ms in 50 pA steps from −400 pA to +500 pA and the number of action potentials were counted at each step. Current injections were conducted prior to (baseline) and after a 10-15 min wash on of the D1 receptor agonist SKF 81297 (10 μM). For depolarization-induced suppression of excitation (DSE), EPSCs were evoked at a rate of 0.2 Hz for a 30 sec baseline followed by a 10 sec depolarization to +50 mV. After depolarization, EPSCs were monitored for 120 sec at 0.2 Hz. Data were averaged over 3 trials for each neuron. EPSC amplitudes were normalized to the average of the baseline period and the magnitude of DSE was measured as the first point after depolarization. Data from each neuron within a treatment group was combined and presented as mean ± SEM.

### Rotational behavior

To probe for changes in D1 receptor function *in vivo*, mice were tested over a three-week period following the conclusion CIE exposure. After cannula guided injections, mice were placed in a round container and Bonsai software (Lopes et al., 2015) was used for video recording and tracking orientation (radians). The number of rotations was counted using custom code in MATLAB (Mathworks Inc., Natick, MA) in which the difference of orientation at each frame was added until the sum was greater than or equal to 6.28 radians or one full clockwise rotation. The sum total was reset to 0 with the count of each rotation. Counterclockwise turns were counted when the sum was less than or equal to −6.28 radians. Each mouse was tested once for one hour then immediately sacrificed for brain extraction and confirmation of cannula placement.

### Immunoblotting

Tissue punches from the medial DS were prepared using a disposable biopsy puncher (Integra, #3331-A) from fresh frozen brains of CIE or air exposed mice and kept at −80°C until further processing. Proteins were extracted in ice-cold lysis buffer containing 150mM NaCl, 50mM Tris-HCl, 1% Triton X-100, pH 7.4 and supplemented with protease inhibitors (Sigma, Complete Mini Protease Inhibitor Cocktail, #11836153001) using a probe tip sonicator (Sonics, Vibra Cell). Homogenates were further incubated on a rotator for 1h at 4°C and centrifuged at 12,000 x g for 15min at 4°C. Supernatant was kept and protein concentration was measured using the BCA method (Pierce BCA Protein Assay, Thermo Fisher, #23225). Samples were incubated in 2x Laemmli Buffer (120mM Tris-HCl, pH=6.8, 20% (w/v) glycerol, 4% SDS, 10% 2-mercaptoethanol, 0.02% bromphenol blue) for 15min at 50°C and 5μg of protein were separated on a 4-20% SDS PAGE (Biorad, #4561096) and blotted onto nitrocellulose membranes by wet transfer. Membranes were stained with Ponceau S solution to determine equal protein loading before blocking in 0.1% PBST (phosphate-buffered saline, 0.1% Tween-20) containing 5% non-fat dry milk for 1h at room temperature. Incubation with primary antibodies (guinea pig Anti-D1 R 1:1000, Frontier Institute, #2571594; rabbit Anti-D2R 1:1000; Frontier Institute, #2571596; rabbit Anticofilin [D3F9] 1:10,000, Cell Signaling Technology, #5175) in blocking buffer was performed overnight at 4°C. Next day, membranes were washed three times in 0.1% PBST and incubated with secondary antibodies (donkey Anti-guinea pig-Alexa-488 and/or donkey Anti-rabbit-Alexa-647; 1:1000; Jackson Immuno Research) in blocking solution for two hours at room temperature before another three 15min washes in 0.1% PBST. Membranes were briefly rinsed in PBS before imaging using the Biorad ChemiDoc MP imaging system. Band densities (bands at ca. 75-100kD for D1R or D2R) were quantified using Image J (NIH), normalized to cofilin (loading control; ca. 19kD) and are expressed as relative intensities.

### Cannula guided drug injections

Mice were implanted with bilateral cannula (Plastics One) targeting the medial DS ([A], 0.5; [M], ± 1.5; [V], 3.25) under stereotaxic guidance and were given one week of recovery prior to CIE procedures. For rotation behaviors, mice received a unilateral injection (left hemisphere) of 300 nL of SKF81297 (13.49 mM) at a rate of 100 nL/min 20 minutes prior to testing. For CB1 receptor antagonist effects on devaluation, 300nL of an 8 μM solution of SR141716A made up of 75% DMSO and 25% saline was injected into each hemisphere 30 minutes prior to prefeeding for outcome devaluation testing.

### Statistical analysis

Statistical significance was defined as an alpha of p < 0.05. Statistical analysis was performed using GraphPad Prism 6 (GraphPad Software) and MatLab. To determine if there were significant differences in photometry data between groups, permutation testing was performed (Jean-Richard-dit-Bressel et al., 2020; Pascoli et al., 2018) in which trials of event related activity from each treatment group were combined and randomly partitioned 1000 times to compute Z-score means and standard deviations across 50 ms bins. To account for the potential false-discovery rate associated with doing a high number of tests, we applied the Benjamini and Hochberg False Discovery Rate correction (Benjamini & Hochberg, 1995). We also put a significance threshold that required at least 4 consecutive time points meet the threshold of p < 0.05. Behavioral and electrophysiology data were analyzed using one-way ANOVA, repeated measures ANOVA with pre-planned Bonferroni-corrected post hoc analyses or a one or two-tailed t-test.

## Author contributions

Rafael Renteria and Christina M Gremel conceptualized and designed the experiments. Cristian Cazares wrote Matlab scripts for analysis of photometry and behavioral data. Rafael Renteria, Emily T Baltz, and Drew C Schreiner performed surgeries and behavioral experiments. Ege A Yalcinbas and Emily T Baltz wrote Matlab scripts for analysis of behavioral data. Thomas Steinkellner and Thomas S Hnasko contributed to and oversaw the immunoblotting experiments. Rafael Renteria performed electrophysiological recordings and analysis. Rafael Renteria and Christina M Gremel wrote the manuscript, with all others contributing to the editing.

---

**Figure 1-figure supplement 1.**
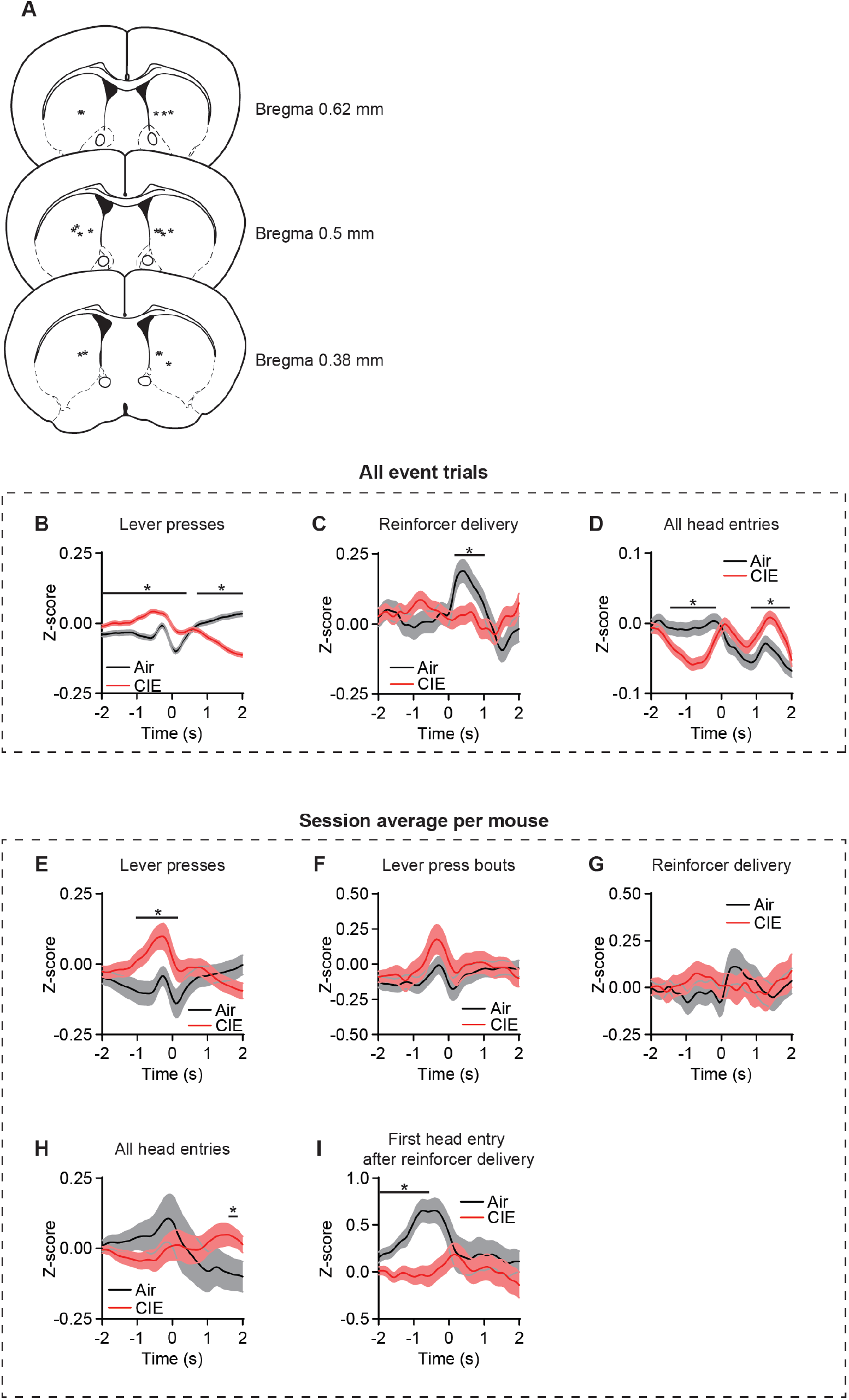
**(A)** Schematic representation of fiber placement in DS. **(B)** Z-score of ΔF/F GCaMP6s trace including all event trials during lever presses (two-sided FDR-corrected permutation test, Air: *n* = 12673; CIE: *n* = 14871), **(C)** reinforcer delivery (two-sided FDR-corrected permutation test, Air: *n* = 569; CIE: *n* = 705), and **(D)** all head entries (two-sided FDR-corrected permutation test, Air: *n* = 8583; CIE: *n* = 8967). **(E)** Z-score of ΔF/F GCaMP6s trace including the session average for each mouse during lever presses (two-sided FDR-corrected permutation test, Air: *n* = 36; CIE: *n* = 34) **(F)** lever press bouts (two-sided FDR-corrected permutation test, Air: *n* = 35; CIE: *n* = 36), **(G)** reinforcer delivery (two-sided permutation test, Air: *n* = 32; CIE: *n* = 36), **(H)** all head entries (two-sided FDR-corrected permutation test, Air: *n* = 34; CIE: *n* = 31), and **(I)** the first head entry after reinforcer delivery (two-sided FDR-corrected permutation test, Air: *n* = 34; CIE: *n* = 38). Two-sided FDR-corrected permutation test, *p < 0.05

**Figure 2-figure supplement 2.**
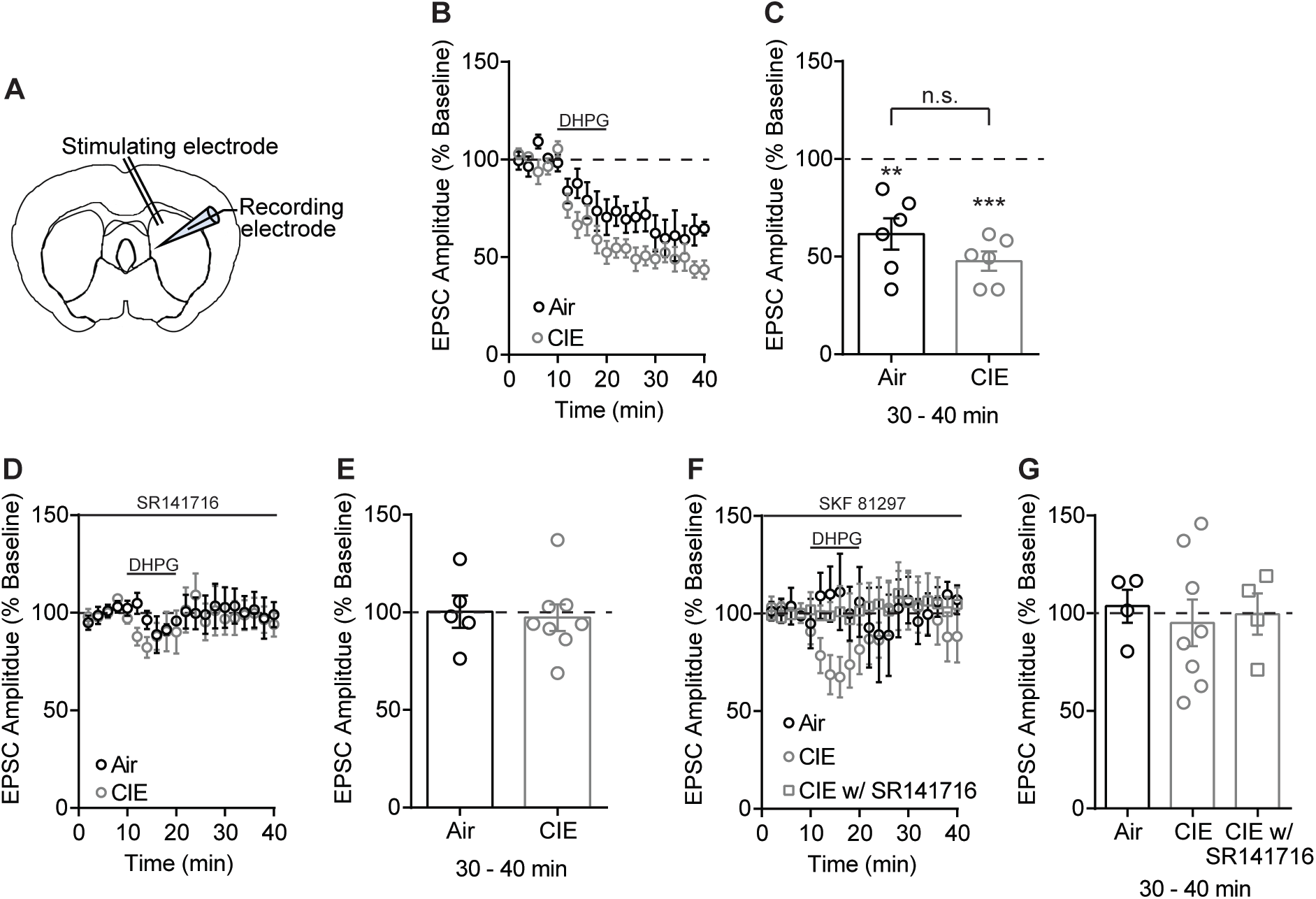
No difference between Air and CIE in the expression of DHPG-LTD of electrically evoked EPSCs. **(A)** Schematic of local electrical stimulation and recording site in the DS. **(B)** DHPG-LTD of electrically evoked EPSCs expressed in both Air and CIE mice. **(C)** Bar graph representing the percentage change from baseline (min 0–10) after bath application of DHPG (min 30–40) (Air: 61.62 ± 8.03% of baseline at 30-40 min, *n* = 6, paired t test vs. baseline: t_5_ = 4.79, p<0.01; CIE: 47.69 ± 4.95% of baseline at 30-40 min, *n* = 6, paired t test vs. baseline: t_5_ = 10.57, p = 0.0001; unpaired t test Air vs. CIE: t_10_ = 1.48, p = 0.17) **(D)** DHPG-LTD is blocked by a CB1 receptor antagonist, SR141716 (1 μM). **(E)** Bar graphs representing the percentage change from baseline after bath application of DHPG in the presence of SR141716 (Air: 100.3 ± 8.27% of baseline at 30-40 min, *n*=5, paired t test vs. baseline: t_4_ = 0.04, p = 0.97; CIE: 97.25 ± 6.87% of baseline at 30-40 min, *n* = 8, paired t test vs. baseline: t_7_ = 0.40, p = 0.70). **(F)** D1 agonist (3 μM SKF 81297) blocks the expression of DHPG-LTD in both Air and CIE. **(G)** Bar graphs representing the percentage change from baseline after bath application of DHPG in the presence of SKF 81297 or SKF 81297 with SR141716 (Air: 103.6 ± 8.45% of baseline at 30-40 min, *n* = 4, paired t test vs. baseline: t_3_ = 0.43, p = 0.70; CIE: 95.11 ± 11.97% of baseline at 30-40 min, *n* = 8, paired t test vs. baseline: t_7_ = 0.41, p = 0.70). Short term depression in CIE mice (74.04 ± 9.88% of baseline at 10-20 min, *n* = 8, paired t test vs. baseline: t_7_ = 2.63, p = 0.03) is CB1 receptor dependent (107.10 ± 10.13% of baseline at 10-20 min, *n* = 4, paired t test vs. baseline: t_3_ = 0.70, p = 0.53; 99.61 ± 10.58% of baseline at 30-40 min, *n* = 4, paired t test vs. baseline: t_3_ = 0.04, p = 0.97). Data points and bar graphs represent the mean ± SEM.

**Figure 3-figure supplement 3.**
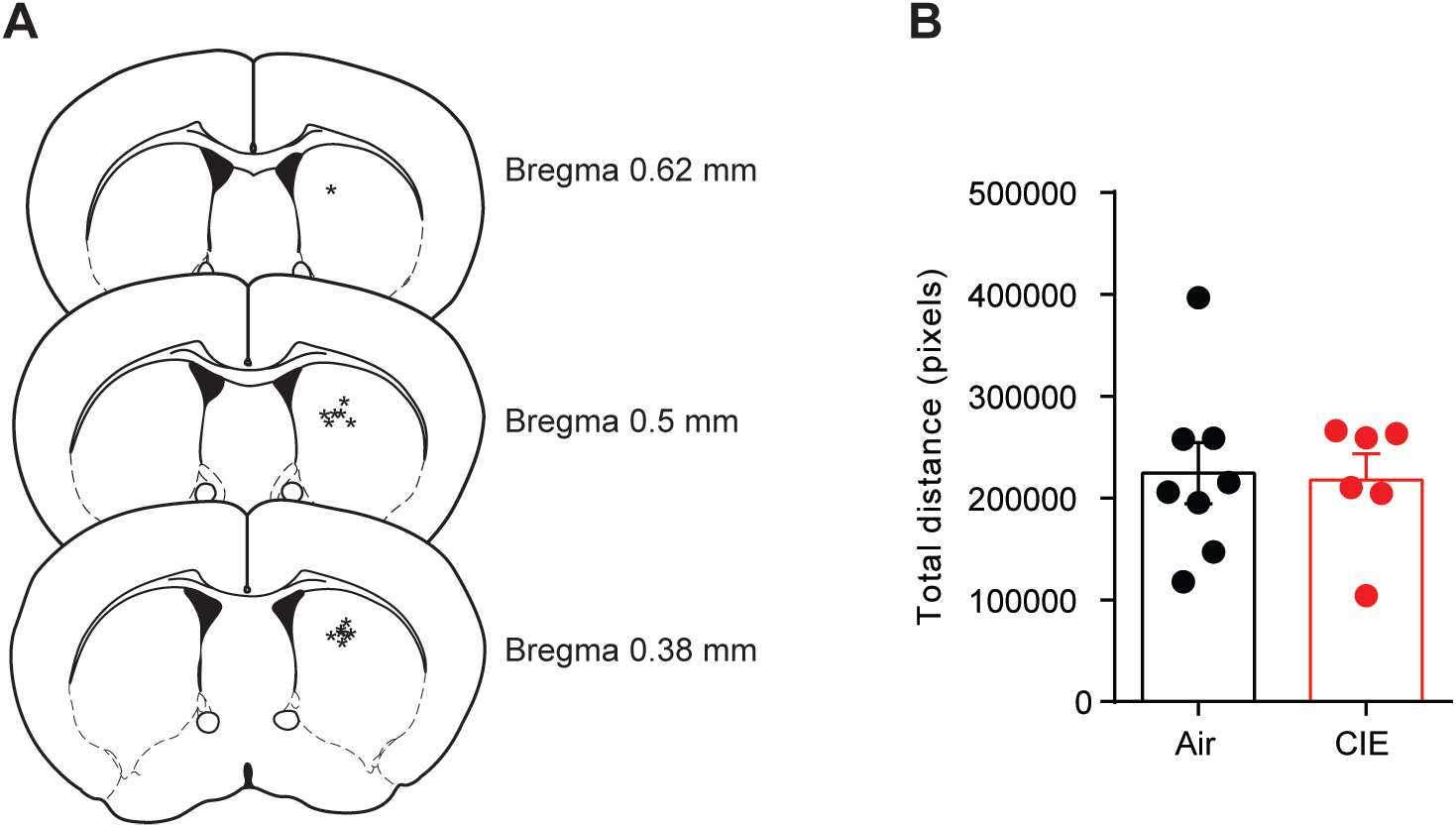
Unilateral D1 agonist injection and distance traveled. **(A)** Schematic representation of cannula tip placement in DS for unilateral injections. **(B)** Total distance traveled in Air and CIE mice. Bar graphs represent the mean ± SEM.

**Figure 3-figure supplement 4.**
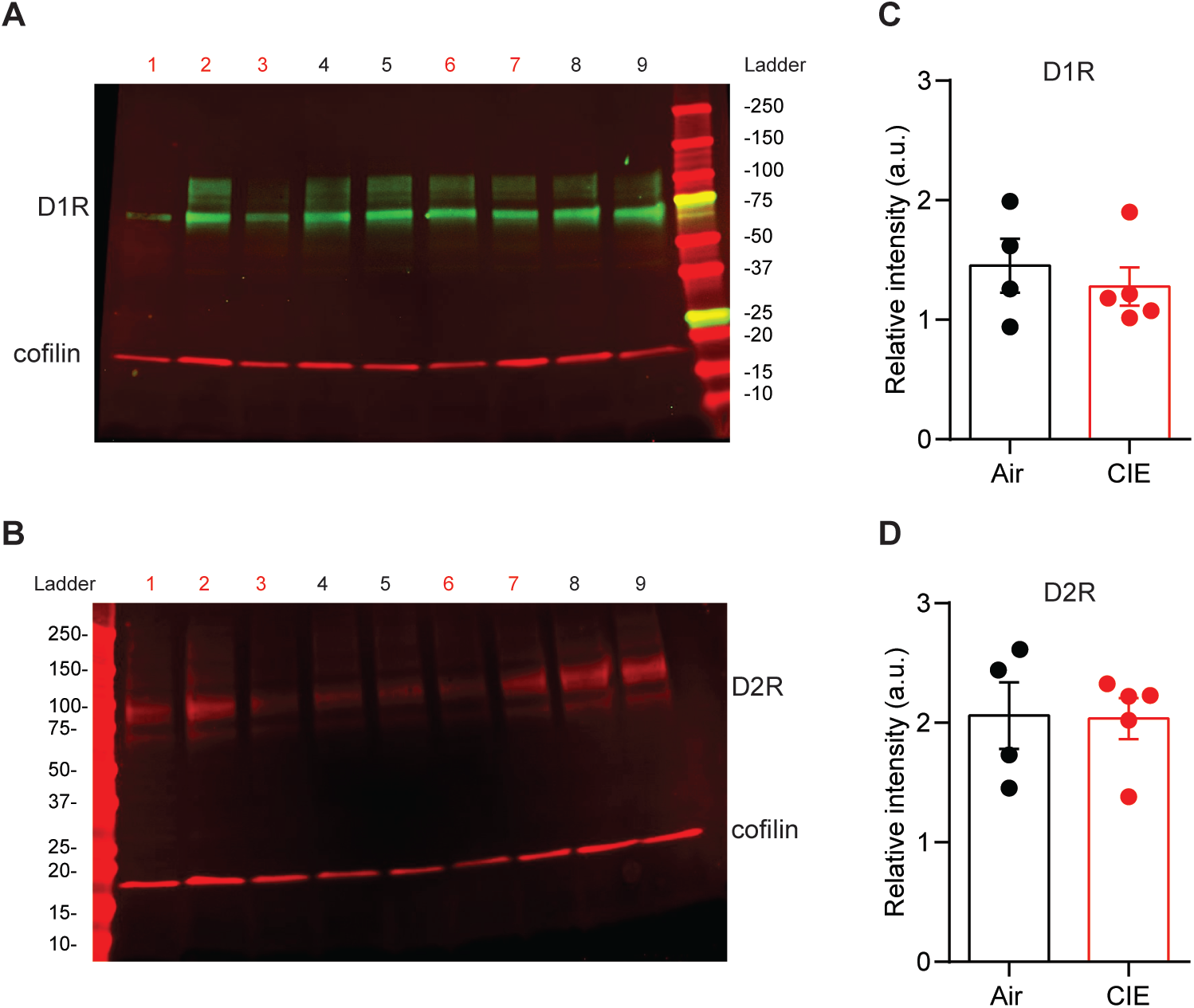
No change in DS D1 or D2 receptor expression in CIE mice compared to Air controls. **(A)** Example immunoblots for D1 and **(B)** D2 dopamine receptors (Air = 4, 5, 8, 9; CIE = 1, 2, 3, 6, 7). **(C)** Band density for D1 and **(D)** D2 receptors normalized to coflin averaged across two runs. Bar graphs represent the mean ± SEM.

**Figure 4-figure supplement 5.**
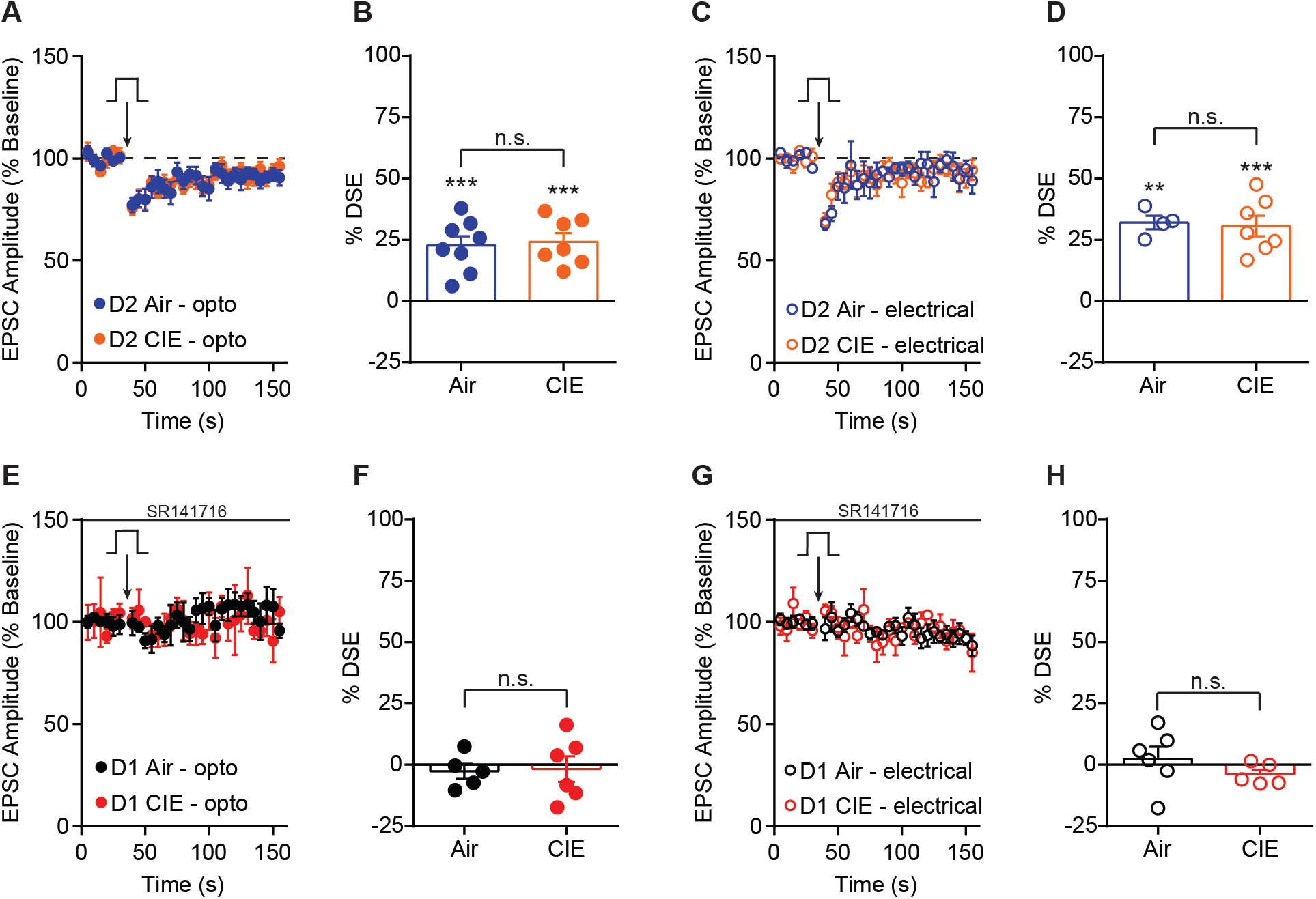
DSE in D2 SPNs and effect of SR141716 on DSE in D1 SPNs. **(A)** Depolarization induced suppression of excitation (DSE) of OFC inputs to D2 SPNs in Air control and CIE mice. **(B)** Bar graphs representing the percentage change from baseline immediately after depolarization. **(C)** DSE of excitatory inputs to D2 SPNs using electrical stimulation in Air and CIE mice. **(D)** Bar graphs representing the percentage change from baseline immediately after depolarization. **(E)** DSE of OFC inputs to D1 SPNs in the presence of a CB1 receptor antagonist, SR141716 (1 μM) in Air and CIE mice. **(F)** Bar graphs representing the percentage change from baseline immediately after depolarization. **(G)** DSE of excitatory inputs to D1 SPNs using electrical stimulation in the presence of a CB1 receptor antagonist, SR141716 (1 μM) in Air and CIE mice. **(H)** Bar graphs representing the percentage change from baseline immediately after depolarization. Data points and bar graphs represent the mean ± SEM. Paired (vs. baseline) two-tailed t-test, **p < 0.01, ***p < 0.001.

**Figure 4-figure supplement 6.**
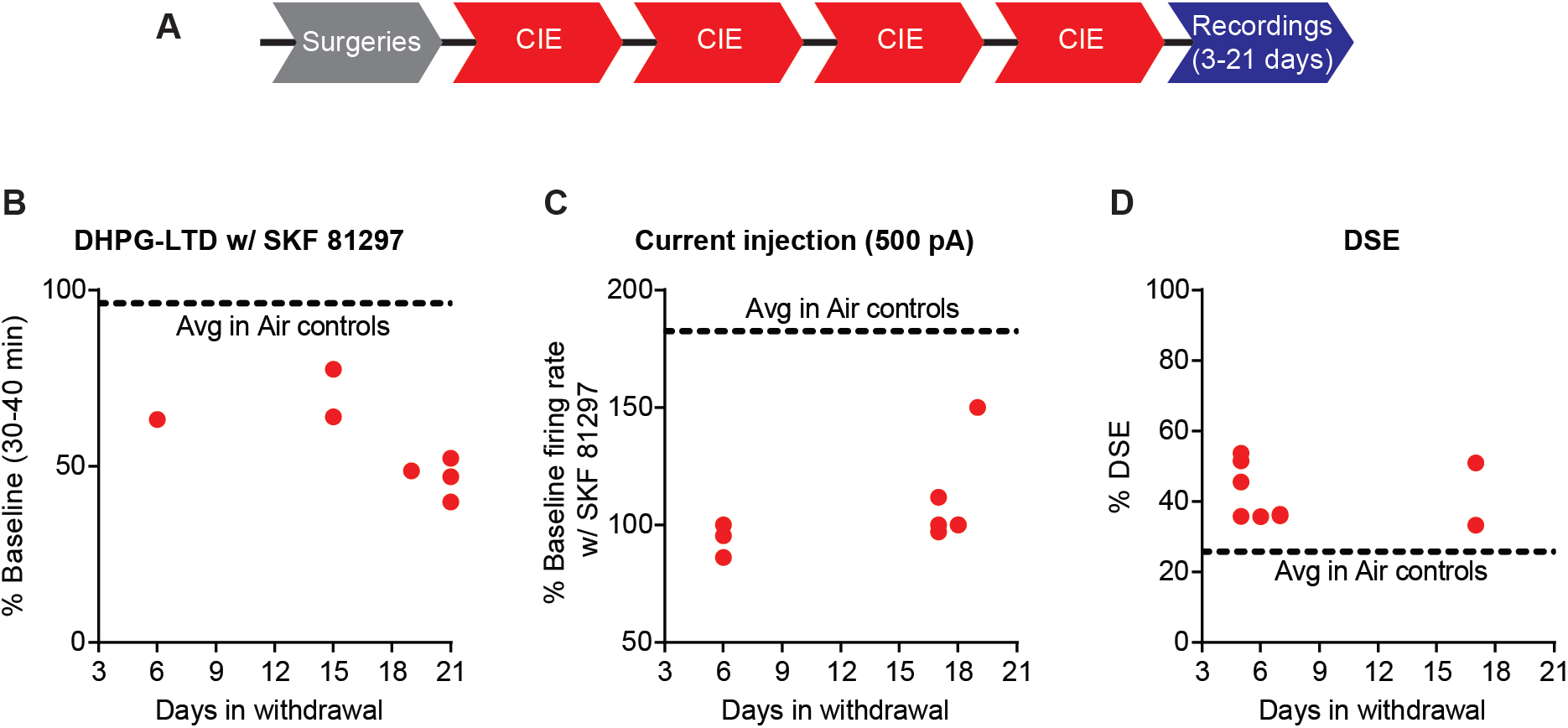
CIE induced effects on D1 receptor function and DSE across recording days. **(A)** Experimental timeline that includes surgeries followed by 4 cycles of CIE exposure and whole-cell recordings 3-21 days in withdrawal. **(B)** Expression of DHPG-LTD in the presence of SKF 81297 was persistent across the withdrawal period. **(C)** SKF 81297 had no effect on D1 SPN firing of CIE mice across the withdrawal period. **(D)** DSE was increased in CIE mice across the withdrawal period.

**Figure 5-figure supplement 7.**
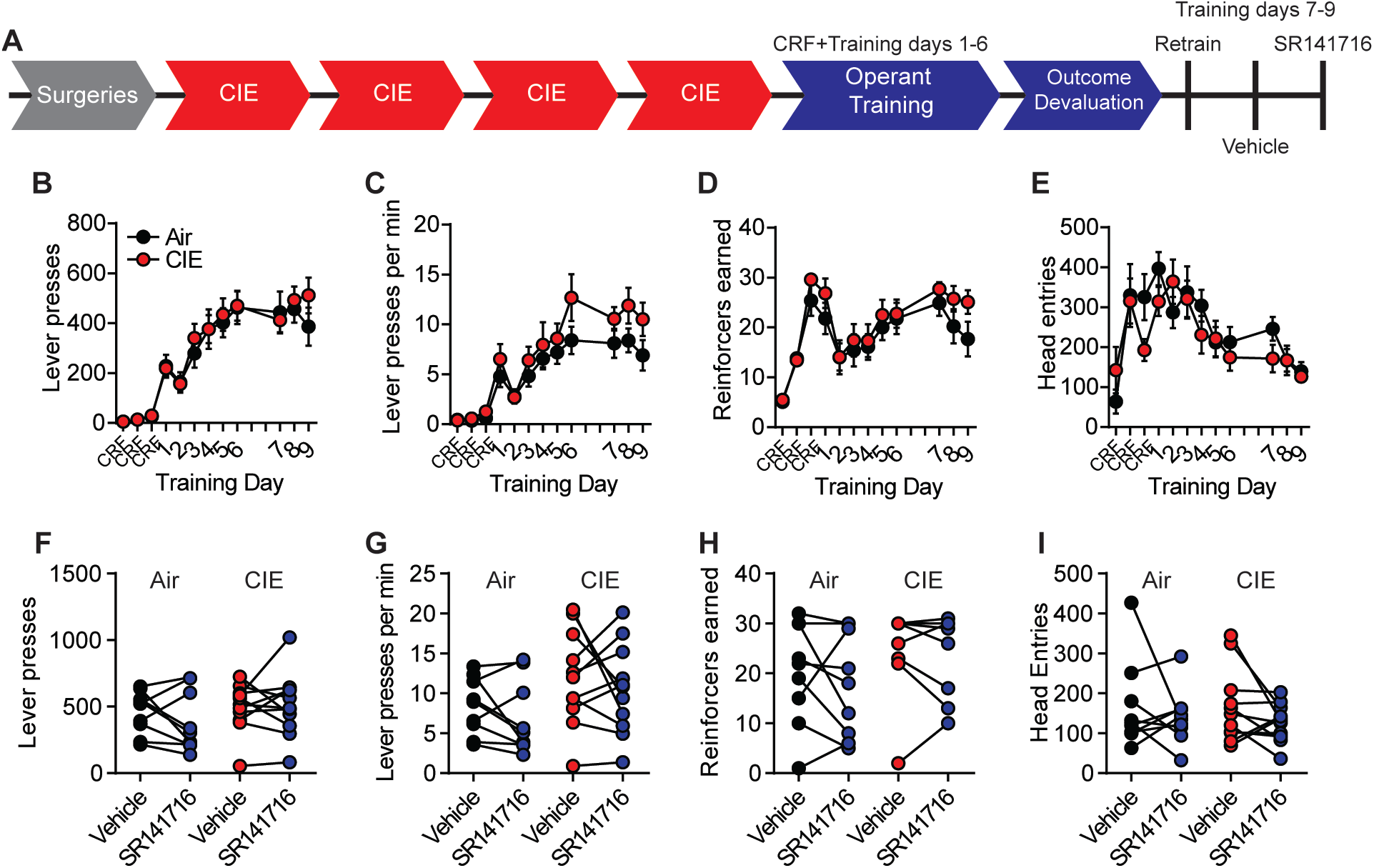
CB1 antagonist did not alter lever press behavior between training days. **(A)** Experimental timeline that includes training after outcome devaluation and injections with vehicle or SR141716. **(B)** Lever presses, **(C)** response rate, **(D)** reinforcers earned, and **(E)** head entries across all training days and retraining and testing days following outcome devaluation. **(F)** The number of lever presses, **(G)** response rate, **(H)** reinforcers earned, **(I)** head entries during vehicle or SR141716 injection test sessions in Air and CIE mice. Data points across training days represent the mean ± SEM.

**Figure 5-figure supplement 8.**
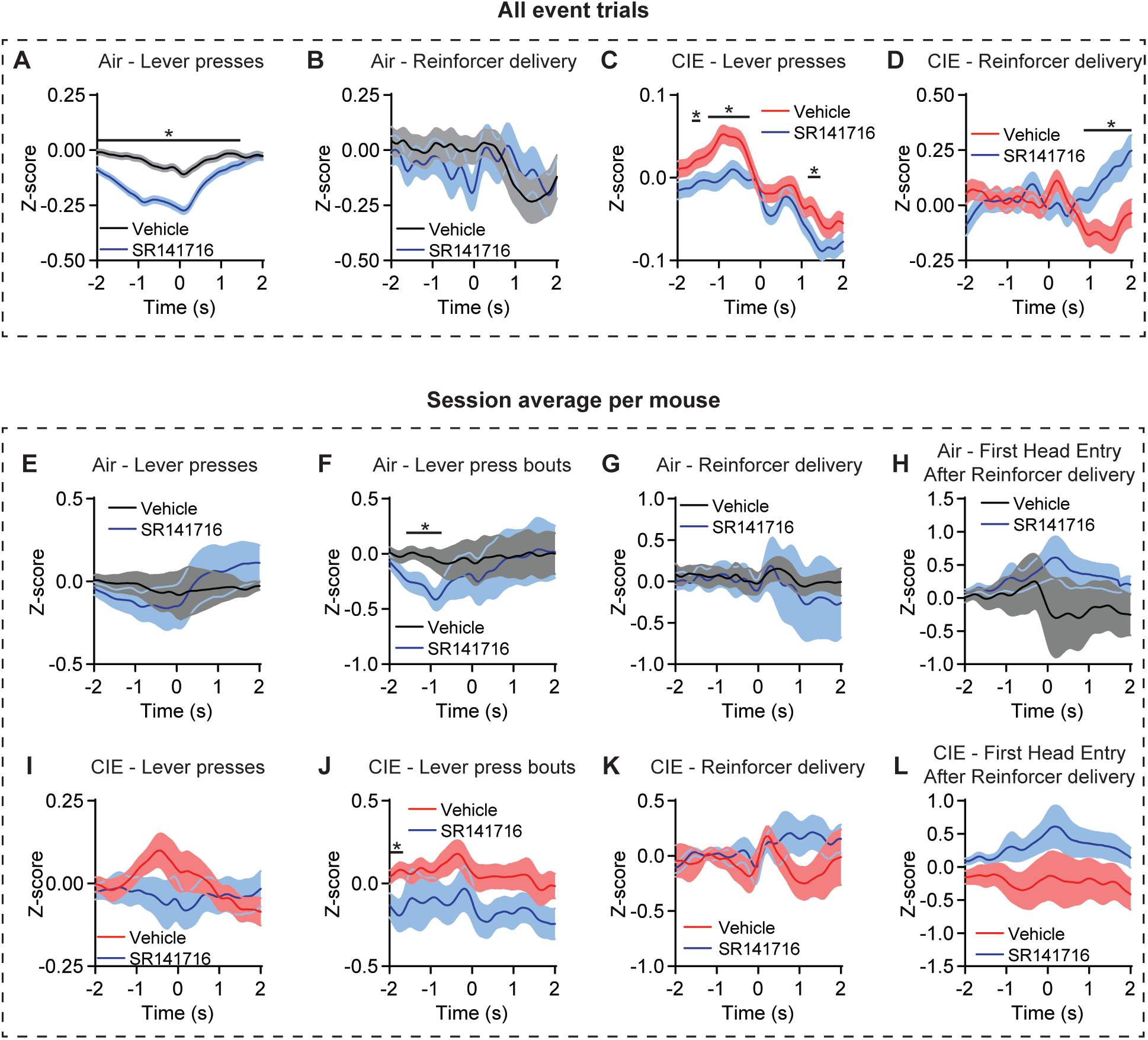
OFC terminal activity in DS with vehicle or CB1 antagonist. **(A)** Z-score of ΔF/F GCaMP6s trace including all event trials in vehicle or SR141716 test sessions during lever presses (two-sided FDR-corrected permutation test, Vehicle: *n* = 3858; SR141716: *n* = 3773) and **(B)** reinforcer delivery (two-sided FDR-corrected permutation test, Vehicle: *n* = 177; SR141716: *n* = 160) in Air mice. **(C)** Z-score of ΔF/F GCaMP6s trace including all event trials in vehicle or SR141716 test sessions during lever presses (two-sided FDR-corrected permutation test, Vehicle: *n* = 5420; SR141716: *n* = 4721) and **(D)** reinforcer delivery (two-sided FDR-corrected permutation test, Vehicle: *n* = 274; SR141716: *n* = 217) in CIE mice. **(E)** Z-score of ΔF/F GCaMP6s trace including the session average for each Air mouse in vehicle or SR141716 test sessions during lever presses, (two-sided FDR-corrected permutation test, Vehicle: *n* = 8; SR141716: *n* = 10), **(F)** lever press bouts (two-sided FDR-corrected permutation test, Vehicle: *n* = 8; SR141716: *n* = 9), **(G)** reinforcer delivery (two-sided FDR-corrected permutation test, Vehicle: *n* = 8; SR141716: *n* = 10), and **(H)** the first head entry after reinforcer delivery (two-sided FDR-corrected permutation test, Vehicle: *n* = 8; SR141716: *n* = 10). **(I)** Z-score of ΔF/F GCaMP6s trace including the session average for each CIE mouse in vehicle or SR141716 test sessions during lever presses, (two-sided FDR-corrected permutation test, Vehicle: *n* = 11; SR141716: *n* = 10), **(J)** lever press bouts (two-sided FDR-corrected permutation test, Vehicle: *n* = 9; SR141716: *n* = 10), **(K)** reinforcer delivery (two-sided FDR-corrected permutation test, Vehicle: *n* = 12; SR141716: *n* = 9), and **(L)** the first head entry after reinforcer delivery (two-sided FDR-corrected permutation test, Vehicle: *n* = 12; SR141716: *n* = 10). Two-sided permutation test *p < 0.05.

**Figre 6-Supplemental Figure 9.**
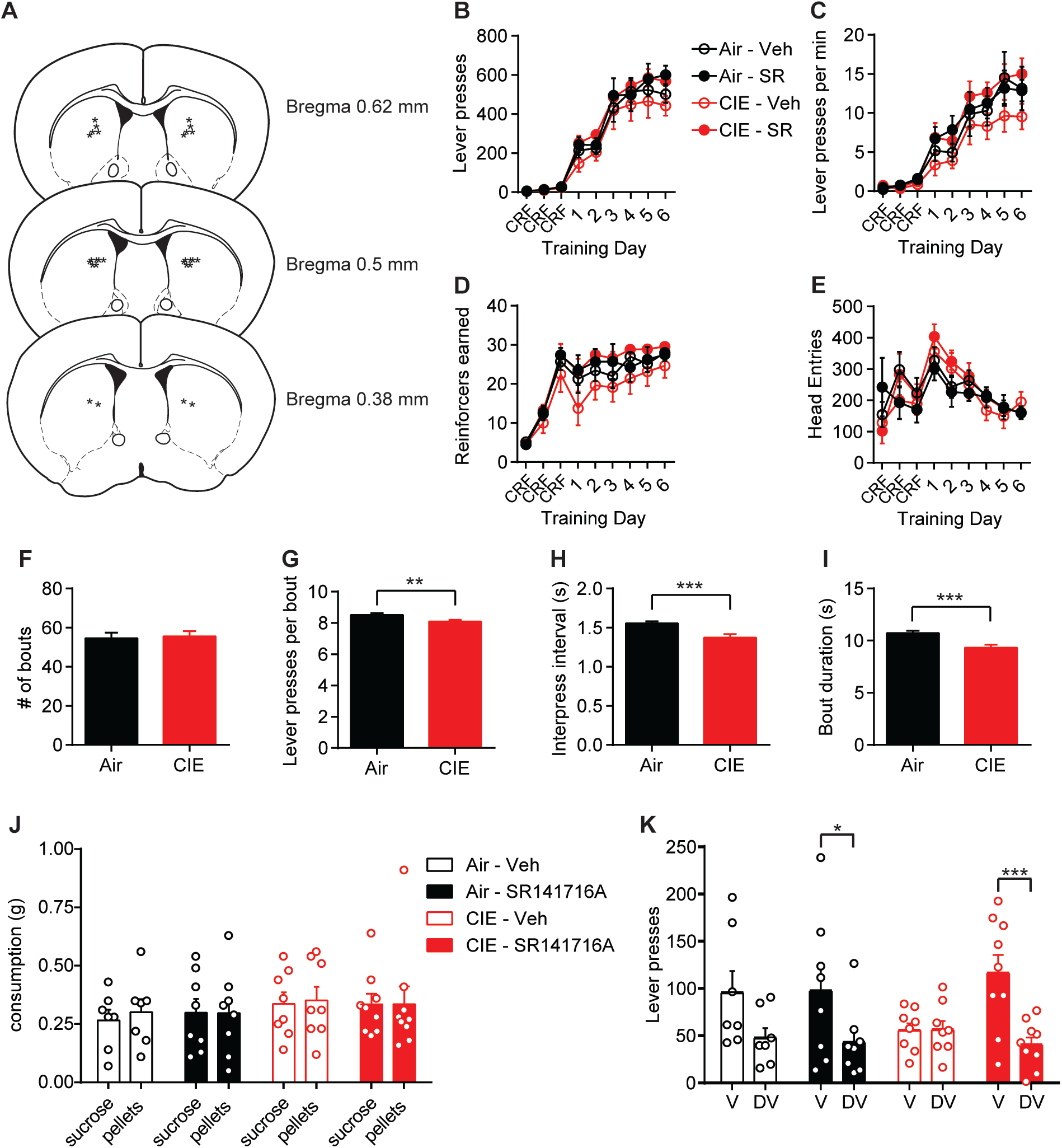
Behavioral data during lever press training and outcome devaluation in treatment and control groups of Air and CIE mice. **(A)** Schematic representation of bilateral cannula placement in DS. **(B)** Lever presses across training days for each treatment group (repeated-measures ANOVA: main effect of Time: F_(8, 224)_ = 157.0, p < 0.0001); no main effect of Treatment or interaction, Fs < 1.08). **(C)** Response rate of lever pressing during acquisition (repeated-measures ANOVA: main effect of Time: F(8, 224) = 86.80, p < 0.0001); no main effect of Treatment or interaction, Fs < 1.10). **(D)** Reinforcers earned across training days (repeated-measures ANOVA: main effect of Time: F_(8, 224)_ = 70.03, p < 0.0001); no main effect of Treatment or interaction, Fs < 1.40). **(E)** Head entries across training days (repeated-measures ANOVA: main effect of Time: F_(8, 224)_ = 10.68, p < 0.0001); no main effect of Treatment or interaction, Fs < 1.02). **(F)** Average number of lever press bouts (Air: 54.58 ± 2.83, *n* = 60; CIE: 55.54 ± 2.61, *n* = 68; unpaired t test: t_126_ = 0.25, p = 0.80), **(G)** lever presses per bout (Air: 8.52 ± 0.12, *n* = 3275; CIE: 8.10 ± 0.10, *n* = 3777; unpaired t test: t_7050_ = 2.72, p < 0.01), **(H)** interpress interval per lever press bout (Air: 1.56 ± 0.03, *n* = 3275; CIE: 1.37 ± 0.05, *n* = 3777; unpaired t test: t7050 = 3.33, p < 0.001), and **(I)** bout duration in Air and CIE mice (Air: 10.72 ± 0.23, *n* = 3275; CIE: 9.34 ± 0.27, *n* = 3777; unpaired t test: t7050 = 3.85, p < 0.001). **(J)** Consumption of sucrose and pellets on valued and devalued days respectively (repeated-measures ANOVA: no interaction or main effects of Time or Treatment, Fs < 0.41). **(K)** Lever presses on Valued and Devalued days for each treatment group (repeated-measures ANOVA: Valuation state x Treatment interaction: F_(3, 28)_ = 3.28, p < 0.05; main effect of Valuation state: F(1, 28) = 23.71, p < 0.0001); no main effect of Treatment F < 0.49). Data points and bar graphs represent the mean ± SEM. Bonferroni corrected post-hoc test of repeated-measures ANOVA or unpaired two-tailed t-test, *p < 0.05, **p < 0.01, ***p < 0.001.

## References

Abrahao KP, Salinas AG, Lovinger DM. 2017. Alcohol and the Brain: Neuronal Molecular Targets, Synapses, and Circuits. Neuron 96:1223–1238. doi:10.1016/j.neuron.2017.10.032

Adermark L, Lovinger DM. 2009. Frequency-dependent inversion of net striatal output by endocannabinoid-dependent plasticity at different synaptic inputs. J Neurosci 29:1375–1380. doi:10.1523/JNEUROSCI.3842-08.2009

Adermark L, Talani G, Lovinger DM. 2009. Endocannabinoid-dependent plasticity at GABAergic and glutamatergic synapses in the striatum is regulated by synaptic activity. Eur J Neurosci 29:32–41. doi:10.1111/j.1460-9568.2008.06551.x

Ahmari SE, Spellman T, Douglass NL, Kheirbek MA, Simpson HB, Deisseroth K, Gordon JA, Hen R. 2013. Repeated cortico-striatal stimulation generates persistent OCD-like behavior. Science (80-) 340:1234–9. doi:10.1126/science.1234733

American Psychiatric Association. 2013. Diagnostic and Statistical Manual of Mental Disorders (DSM-5), Arlington, VA. doi:10.1176/appi.books.9780890425596.893619

André VM, Cepeda C, Cummings DM, Jocoy EL, Fisher YE, William Yang X, Levine MS. 2010. Dopamine modulation of excitatory currents in the striatum is dictated by the expression of D1 or D2 receptors and modified by endocannabinoids. Eur J Neurosci 31:14–28. doi:10.1111/j.1460-9568.2009.07047.x

Becker HC. 1994. Positive relationship between the number of prior ethanol withdrawal episodes and the severity of subsequent withdrawal seizures. Psychopharmacology (Berl) 116:26–32.

Becker HC, Lopez MF. 2004. Increased ethanol drinking after repeated chronic ethanol exposure and withdrawal experience in C57BL/6 mice. Alcohol Clin Exp Res 28:1829–1838.

Belin D, Belin-Rauscent A, Murray JE, Everitt BJ. 2013. Addiction: Failure of control over maladaptive incentive habits, Current Opinion in Neurobiology. doi:10.1016/j.conb.2013.01.025

Benjamini Y., Hochberg Y. 1995. Controlling the false discovery rate: a practical and powerful approach to multiple testing. J Royal Statistical Society Series B 57(1): 289–300.

Boettiger C a, Mitchell JM, Tavares VC, Robertson M, Joslyn G, D’Esposito M, Fields HL. 2007. Immediate reward bias in humans: fronto-parietal networks and a role for the catechol-O-methyltransferase 158(Val/Val) genotype. J Neurosci 27:14383–91. doi:10.1523/JNEUROSCI.2551-07.2007

Castillo PE, Younts TJ, Chávez AE, Hashimotodani Y. 2012. Endocannabinoid Signaling and Synaptic Function. Neuron. doi:10.1016/j.neuron.2012.09.020

Catafau AM, Etcheberrigaray A, Perez de los Cobos J, Estorch M, Guardia J, Flotats A, Bernà L, Marí C, Casas M, Carrió I. 1999. Regional cerebral blood flow changes in chronic alcoholic patients induced by naltrexone challenge during detoxification. J Nucl Med 40:19–24.

Christian DT, Alexander NJ, Diaz MR, Robinson S, McCool BA. 2012. Chronic intermittent ethanol and withdrawal differentially modulate basolateral amygdala AMPA-type glutamate receptor function and trafficking. Neuropharmacology 62:2429–2438. doi:10.1016/j.neuropharm.2012.02.017

Corbit LH, Nie H, Janak PH. 2012. Habitual alcohol seeking: Time course and the contribution of subregions of the dorsal striatum. Biol Psychiatry 72:389–395. doi:10.1016/j.biopsych.2012.02.024

Covey DP, Mateo Y, Sulzer D, Cheer JF, Lovinger DM. 2017. Endocannabinoid modulation of dopamine neurotransmission. Neuropharmacology. doi:10.1016/j.neuropharm.2017.04.033

Cuzon Carlson VC, Seabold GK, Helms CM, Garg N, Odagiri M, Rau AR, Daunais J, Alvarez VA, Lovinger DM, Grant KA. 2011. Synaptic and morphological neuroadaptations in the putamen associated with long-term, relapsing alcohol drinking in primates. Neuropsychopharmacology 36:2513–28. doi:10.1038/npp.2011.140

DePoy L, Daut R, Brigman JL, MacPherson K, Crowley N, Gunduz-Cinar O, Pickens CL, Cinar R, Saksida LM, Kunos G, Lovinger DM, Bussey TJ, Camp MC, Holmes A. 2013. Chronic alcohol produces neuroadaptations to prime dorsal striatal learning. Proc Natl Acad Sci U S A 110:14783–8. doi:10.1073/pnas.1308198110

Diana MA, Marty A. 2004. Endocannabinoid-mediated short-term synaptic plasticity: Depolarization-induced suppression of inhibition (DSI) and depolarization-induced suppression of excitation (DSE). Br J Pharmacol 142:9–19. doi:10.1038/sj.bjp.0705726

Dickinson A, Nicholas DJ, Adams CD. 1983. The effect of the instrumental training contingency on susceptibility to reinforcer devaluation. Q J Exp Psychol Sect B 35:35–51. doi:10.1080/14640748308400912

Dickinson A, Wood N, Smith JW. 2002. Alcohol seeking by rats: action or habit? Q J Exp Psychol B 55:331–48. doi:10.1080/0272499024400016

Duka T, Trick L, Nikolaou K, Gray MA, Kempton MJ, Williams H, Williams SCR, Critchley HD, Stephens DN. 2011. Unique brain areas associated with abstinence control are damaged in multiply detoxified alcoholics. Biol Psychiatry 70:545–52. doi:10.1016/j.biopsych.2011.04.006

Ernst LH, Plichta MM, Dresler T, Zesewitz AK, Tupak S V., Haeussinger FB, Fischer M, Polak T, Fallgatter AJ, Ehlis AC. 2014. Prefrontal correlates of approach preferences for alcohol stimuli in alcohol dependence. Addict Biol 19:497–508. doi:10.1111/adb.12005

Everitt BJ, Robbins TW. 2016. Drug Addiction: Updating Actions to Habits to Compulsions Ten Years On. Annu Rev Psychol 67:23–50. doi:10.1146/annurev-psych-122414-033457

Everitt BJ, Robbins TW. 2005. Neural systems of reinforcement for drug addiction: from actions to habits to compulsion. Nat Neurosci 8:1481–9. doi:10.1038/nn1579

Fellows LK. 2007. The Role of Orbitofrontal Cortex in Decision Making: A Component Process Account. Ann N Y Acad Sci 421–30. doi:10.1196/annals.1401.023

Forbes EE, Rodriguez EE, Musselman S, Narendran R. 2014. Prefrontal response and frontostriatal functional connectivity to monetary reward in abstinent alcohol-dependent young adults. PLoS One 9:e94640. doi:10.1371/journal.pone.0094640

Gerdeman GL, Ronesi J, Lovinger DM. 2002. Postsynaptic endocannabinoid release is critical to long-term depression in the striatum. Nat Neurosci 5:446–51. doi:10.1038/nn832

Gianessi CA, Groman SM, Thompson SL, Jiang M, van der Stelt M, Taylor JR. 2020. Endocannabinoid contributions to alcohol habits and motivation: Relevance to treatment. Addict Biol 25:e12768. doi:10.1111/adb.12768

Giuffrida A, Parsons LH, Kerr TM, Rodríguez De Fonseca F, Navarro M, Piomelli D. 1999. Dopamine activation of endogenous cannabinoid signaling in dorsal striatum. Nat Neurosci 2:358–363. doi:10.1038/7268

Glick SD, Jerussi TP, Fleisher LN. 1976. Turning in circles: The neuropharmacology of rotation. Life Sci 18:889–896. doi:10.1016/0024-3205(76)90405-7

Gremel CM, Chancey JH, Atwood BK, Luo G, Neve R, Ramakrishnan C, Deisseroth K, Lovinger DM, Costa RM. 2016. Endocannabinoid Modulation of Orbitostriatal Circuits Gates Habit Formation. Neuron 90:1312–1324. doi:10.1016/j.neuron.2016.04.043

Gremel CM, Costa RM. 2013. Orbitofrontal and striatal circuits dynamically encode the shift between goal-directed and habitual actions. Nat Commun 4:2264. doi:10.1038/ncomms3264

Gremel CM, Lovinger DM. 2017. Associative and sensorimotor cortico-basal ganglia circuit roles in effects of abused drugs. Genes, Brain Behav 16:71–85. doi:10.1111/gbb.12309

Griffin 3rd WC, Lopez MF, Becker HC. 2009. Intensity and duration of chronic ethanol exposure is critical for subsequent escalation of voluntary ethanol drinking in mice. Alcohol Clin Exp Res 33:1893–1900. doi:10.1111/j.1530-0277.2009.01027.x

Heifets BD, Castillo PE. 2009. Endocannabinoid signaling and long-term synaptic plasticity. Annu Rev Physiol 71:283–306. doi:10.1146/annurev.physiol.010908.163149

Heilig M, Egli M, Crabbe JC, Becker HC. 2010. Acute withdrawal, protracted abstinence and negative affect in alcoholism: Are they linked? Addict Biol 15: 169–184. doi:10.1111/j.1369-1600.2009.00194.x

Henderson-Redmond AN, Guindon J, Morgan DJ. 2016. Roles for the endocannabinoid system in ethanol-motivated behavior. Prog Neuro-Psychopharmacology Biol Psychiatry 65:330–339. doi:10.1016/j.pnpbp.2015.06.011

Hermann D, Smolka MN, Wrase J, Klein S, Nikitopoulos J, Georgi A, Braus DF, Flor H, Mann K, Heinz A. 2006. Blockade of cue-induced brain activation of abstinent alcoholics by a single administration of amisulpride as measured with fMRI. Alcohol Clin Exp Res 30:1349–54. doi:10.1111/j.1530-0277.2006.00174.x

Hernandez-Lopez S, Bargas J, Surmeier DJ, Reyes A, Galarraga E. 1997. D1 receptor activation enhances evoked discharge in neostriatal medium spiny neurons by modulating an L-type Ca2+ conductance. J Neurosci 17:3334–3342. doi:10.1016/S0079-6123(08)61352-7

Hilário MRF, Clouse E, Yin HH, Costa RM. 2007. Endocannabinoid signaling is critical for habit formation. Front Integr Neurosci 1:6. doi:10.3389/neuro.07.006.2007

Hirth N, Meinhardt MW, Noori HR, Salgado H, Torres-Ramirez O, Uhrig S, Broccoli L, Vengeliene V, Roßmanith M, Perreau-Lenz S, Köhr G, Sommer WH, Spanagel R, Hansson AC. 2016. Convergent evidence from alcohol-dependent humans and rats for a hyperdopaminergic state in protracted abstinence. Proc Natl Acad Sci 113:3024–3029. doi:10.1073/pnas.1506012113

Hogarth L, Balleine BW, Corbit LH, Killcross S. 2013. Associative learning mechanisms underpinning the transition from recreational drug use to addiction. Ann N Y Acad Sci 1282:12–24. doi:10.1111/j.1749-6632.2012.06768.x

Jean-Richard-dit-Bressel P, Clifford CWG, McNally GP. 2020. Analyzing Event-Related Transients: Confidence Intervals, Permutation Tests, and Consecutive Thresholds. Front Mol Neurosci. doi:10.3389/fnmol.2020.00014

Kano M, Ohno-Shosaku T, Hashimotodani Y, Uchigashima M, Watanabe M. 2009. Endocannabinoid-mediated control of synaptic transmission. Physiol Rev 89:309–80. doi:10.1152/physrev.00019.2008

Kreitzer AC, Malenka RC. 2007. Endocannabinoid-mediated rescue of striatal LTD and motor deficits in Parkinson’s disease models. Nature 445:643–647. doi:10.1038/nature05506

Kreitzer AC, Malenka RC. 2005. Dopamine Modulation of State-Dependent Endocannabinoid Release and Long-Term Depression in the Striatum. J Neurosci 25:10537–10545. doi:10.1523/JNEUROSCI.2959-05.2005

Kreitzer AC, Regehr WG. 2001. Retrograde inhibition of presynaptic calcium influx by endogenous cannabinoids at excitatory synapses onto Purkinje cells. Neuron 29:717–27. doi:10.1016/S0896-6273(01)00246-X

Lin JY. 2010. A user’s guide to channelrhodopsin variants: Features, limitations and future developments. Exp Physiol. doi:10.1113/expphysiol.2009.051961

Lopes G, Bonacchi N, FrazÃ£o J, Neto JP, Atallah B V., Soares S, Moreira L, Matias S, Itskov PM, Correia PA, Medina RE, Calcaterra L, Dreosti E, Paton JJ, Kampff AR. 2015. Bonsai: an event-based framework for processing and controlling data streams. Front Neuroinform 9. doi:10.3389/fninf.2015.00007

Lopez MF, Becker HC. 2005. Effect of pattern and number of chronic ethanol exposures on subsequent voluntary ethanol intake in C57BL/6J mice. Psychopharmacol 181:688–696. doi:10.1007/s00213-005-0026-3

Lovinger DM. 2008. Presynaptic modulation by endocannabinoids. Handb Exp Pharmacol 184:435–477. doi:10.1007/978-3-540-74805-2-14

Lovinger DM, Gremel CM 2020. A circuit-based information approach to substance abuse research. Trends in Neurosci doi.org/10.1016/j.tins.2020.10.005

Lovinger DM, Roberto M. 2013. Synaptic effects induced by alcohol. Curr Top Behav Neurosci. doi:10.1007/7854_2011_143

Lüscher C, Robbins TW, Everitt BJ. 2020. The transition to compulsion in addiction. Nat Rev Neurosci 21:247–263. doi:10.1038/s41583-020-0289-z

Markowitz JE, Gillis WF, Beron CC, Neufeld SQ, Robertson K, Bhagat ND, Peterson RE, Peterson E, Hyun M, Linderman SW, Sabatini BL, Datta SR. 2018. The Striatum Organizes 3D Behavior via Moment-to-Moment Action Selection. Cell 174:44–58.e17. doi:10.1016/j.cell.2018.04.019

Martín AB, Fernandez-Espejo E, Ferrer B, Gorriti MA, Bilbao A, Navarro M, Rodriguez De Fonseca F, Moratalla R. 2008. Expression and function of CB1 receptor in the rat striatum: Localization and effects on D1 and D2 dopamine receptor-mediated motor behaviors. Neuropsychopharmacology 33:1667–1679. doi:10.1038/sj.npp.1301558

Mateo Y, Johnson KA, Covey DP, Atwood BK, Wang HL, Zhang S, Gildish I, Cachope R, Bellocchio L, Guzmán M, Morales M, Cheer JF, Lovinger DM. 2017. Endocannabinoid Actions on Cortical Terminals Orchestrate Local Modulation of Dopamine Release in the Nucleus Accumbens. Neuron 96:1112–1126.e5. doi:10.1016/j.neuron.2017.11.012

Milad MR, Rauch SL. 2012. Obsessive-compulsive disorder: Beyond segregated cortico-striatal pathways. Trends Cogn Sci 16:43–51. doi:10.1016/j.tics.2011.11.003

Moorman DE. 2018. The role of the orbitofrontal cortex in alcohol use, abuse, and dependence. Prog Neuro-Psychopharmacology Biol Psychiatry 87:85–107. doi:10.1016/j.pnpbp.2018.01.010

Nazzaro C, Greco B, Cerovic M, Baxter P, Rubino T, Trusel M, Parolaro D, Tkatch T, Benfenati F, Pedarzani P, Tonini R. 2012. SK channel modulation rescues striatal plasticity and control over habit in cannabinoid tolerance. Nat Neurosci 15:284–93. doi:10.1038/nn.3022

Nimitvilai S, Lopez MF, Mulholland PJ, Woodward JJ. 2016. Chronic Intermittent Ethanol Exposure Enhances the Excitability and Synaptic Plasticity of Lateral Orbitofrontal Cortex Neurons and Induces a Tolerance to the Acute Inhibitory Actions of Ethanol. Neuropsychopharmacology 41:1112–27. doi:10.1038/npp.2015.250

Nimitvilai S, Uys JD, Woodward JJ, Randall PK, Ball LE, Williams RW, Jones BC, Lu L, Grant KA, Mulholland PJ. 2017. Orbitofrontal neuroadaptations and cross-species synaptic biomarkers in heavy drinking macaques. J Neurosci 37:3646–3660. doi:10.1523/JNEUROSCI.0133-17.2017

Parsons LH, Hurd YL. 2015. Endocannabinoid signalling in reward and addiction. Nat Rev Neurosci. doi:10.1038/nrn4004

Pascoli V, Hiver A, Van Zessen R, Loureiro M, Achargui R, Harada M, Flakowski J, Lüscher C. 2018. Stochastic synaptic plasticity underlying compulsion in a model of addiction. Nature 564:366–371. doi:10.1038/s41586-018-0789-4

Pascoli V, Terrier J, Hiver A, Lüscher C. 2015. Sufficiency of Mesolimbic Dopamine Neuron Stimulation for the Progression to Addiction. Neuron 88:1054–1066. doi:10.1016/j.neuron.2015.10.017

Patel S. 2003. Differential Regulation of the Endocannabinoids Anandamide and 2-Arachidonylglycerol within the Limbic Forebrain by Dopamine Receptor Activity. J Pharmacol Exp Ther 306:880–888. doi:10.1124/jpet.103.054270

Pauls DL, Abramovitch A, Rauch SL, Geller DA. 2014. Obsessive-compulsive disorder: An integrative genetic and neurobiological perspective. Nat Rev Neurosci 15:410–24. doi:10.1038/nrn3746

Pava MJ, Woodward JJ. 2012. A review of the interactions between alcohol and the endocannabinoid system: implications for alcohol dependence and future directions for research. Alcohol 46:185–204. doi:10.1016/j.alcohol.2012.01.002

Planert H, Berger TK, Silberberg G. 2013. Membrane Properties of Striatal Direct and Indirect Pathway Neurons in Mouse and Rat Slices and Their Modulation by Dopamine. PLoS One 8. doi:10.1371/journal.pone.0057054

Reinhard I, Leménager T, Fauth-Bühler M, Hermann D, Hoffmann S, Heinz A, Kiefer F, Smolka MN, Wellek S, Mann K, Vollstädt-Klein S. 2015. A comparison of region-of-interest measures for extracting whole brain data using survival analysis in alcoholism as an example. J Neurosci Methods 58–64. doi:10.1016/j.jneumeth.2015.01.001

Renteria R, Baltz ET, Gremel CM. 2018. Chronic alcohol exposure disrupts top-down control over basal ganglia action selection to produce habits. Nat Commun 9:211. doi:10.1038/s41467-017-02615-9

Renteria R, Cazares C, Gremel CM. 2020. Habitual Ethanol Seeking and Licking Microstructure of Enhanced Ethanol Self-Administration in Ethanol-Dependent Mice. Alcohol Clin Exp Res 44:880–891. doi:10.1111/acer.14302

Rex EB, Rankin ML, Ariano M a, Sibley DR. 2008. Ethanol regulation of D(1) dopamine receptor signaling is mediated by protein kinase C in an isozyme-specific manner. Neuropsychopharmacology 33:2900–11. doi:10.1038/npp.2008.16

Robbins TW, Vaghi MM, Banca P. 2019. Obsessive-Compulsive Disorder: Puzzles and Prospects. Neuron 102:27–47. doi:10.1016/j.neuron.2019.01.046

Roberto M, Varodayan FP. 2017. Synaptic targets: Chronic alcohol actions. Neuropharmacology. doi:10.1016/j.neuropharm.2017.01.013

Ron D, Barak S. 2016. Molecular mechanisms underlying alcohol-drinking behaviours. Nat Rev Neurosci. doi:10.1038/nrn.2016.85

Ron D, Messing RO. 2013. Signaling Pathways Mediating Alcohol Effects. Curr Top Behav Neurosci 13:87–126. doi:10.1007/7854_2011_161

Schoenbaum G, Chang CY, Lucantonio F, Takahashi YK. 2016. Thinking Outside the Box: Orbitofrontal Cortex, Imagination, and How We Can Treat Addiction. Neuropsychopharmacology 41:2966–2976. doi:10.1038/npp.2016.147

Schoenbaum G, Roesch MR, Stalnaker TA. 2006. Orbitofrontal cortex, decision-making and drug addiction. Trends Neurosci 29:116–124. doi:10.1016/j.tins.2005.12.006

Shen W, Flajolet M, Greengard P, Surmeier DJ. 2008. Dichotomous dopaminergic control of striatal synaptic plasticity. Science (80-) 321:848–851. doi:10.1126/science.1160575

Shu-Jung Hu S, Mackie K. 2015. Distribution of the endocannabinoid system in the central nervous systemHandbook of Experimental Pharmacology. pp. 59–93. doi:10.1007/978-3-319-20825-1_3

Shuen JA, Chen M, Gloss B, Calakos N. 2008. Drd1a-tdTomato BAC transgenic mice for simultaneous visualization of medium spiny neurons in the direct and indirect pathways of the basal ganglia. J Neurosci 28:2681–5. doi:10.1523/JNEUROSCI.5492-07.2008

Stalnaker TA, Cooch NK, Schoenbaum G. 2015. What the orbitofrontal cortex does not do. Nat Neurosci 18:620–627. doi:10.1038/nn.3982

Surmeier DJ, Ding J, Day M, Wang Z, Shen W. 2007. D1 and D2 dopamine-receptor modulation of striatal glutamatergic signaling in striatal medium spiny neurons. Trends Neurosci. doi:10.1016/j.tins.2007.03.008

Tapert SF, Cheung EH, Brown GG, Frank LR, Paulus MP, Schweinsburg AD, Meloy MJ, Brown SA. 2003. Neural response to alcohol stimuli in adolescents with alcohol use disorder. Arch Gen Psychiatry 60:727–35. doi:10.1001/archpsyc.60.7.727

Trusel M, Cavaccini A, Gritti M, Greco B, Saintot PP, Nazzaro C, Cerovic M, Morella I, Brambilla R, Tonini R. 2015. Coordinated Regulation of Synaptic Plasticity at Striatopallidal and Striatonigral Neurons Orchestrates Motor Control. Cell Rep 13:1353–1365. doi:10.1016/j.celrep.2015.10.009

Tye KM, Prakash R, Kim S-Y, Fenno LE, Grosenick L, Zarabi H, Thompson KR, Gradinaru V, Ramakrishnan C, Deisseroth K. 2011. Amygdala circuitry mediating reversible and bidirectional control of anxiety. Nature 471:358–62. doi:10.1038/nature09820

Volkow ND, Fowler JS. 1994. Brain-imaging studies of the combined use of cocaine and alcohol and of the pharmacokinetics of cocaethylene. NIDA Res Monogr 138:41–56.

Volkow ND, Wang GJ, Fowler JS. 1997. Imaging studies of cocaine in the human brain and studies of the cocaine addict. Ann N Y Acad Sci 41–54. doi:10.1111/j.1749-6632.1997.tb46188.x

Wall NR, De La Parra M, Callaway EM, Kreitzer AC. 2013. Differential innervation of direct- and indirect-pathway striatal projection neurons. Neuron 79:347–60. doi:10.1016/j.neuron.2013.05.014

Wallis JD. 2007. Orbitofrontal Cortex and Its Contribution to Decision-Making. Annu Rev Neurosci 30:31–56. doi:10.1146/annurev.neuro.30.051606.094334

Wu YW, Kim JI, Tawfik VL, Lalchandani RR, Scherrer G, Ding JB. 2015. Input- and cell-type-specific endocannabinoid-dependent LTD in the striatum. Cell Rep 10:75–87. doi:10.1016/j.celrep.2014.12.005

Yin HH, Davis MI, Ronesi J a, Lovinger DM. 2006. The role of protein synthesis in striatal long-term depression. J Neurosci 26:11811–11820. doi:26/46/11811 [pii]\r10.1523/JNEUROSCI.3196-06.2006

